# A genotype-first approach identifies variants for orofacial clefts and other phenotypes in dogs

**DOI:** 10.1101/2024.10.04.616687

**Authors:** Reuben M Buckley, Nüket Bilgen, Alexander C Harris, Peter Savolainen, Cafer Tepeli, Metin Erdoğan, Aitor Serres Armero, Dayna L Dreger, Frank G van Steenbeek, Marjo K Hytönen, Jessica Hale, Hannes Lohi, Bengi Çınar Kul, Adam R Boyko, Elaine A Ostrander

## Abstract

Dog breeding promotes within-group homogeneity through conformation to strict breed standards, and also drives between-group heterogeneity in pursuit of characteristic breed traits. There are over 350 recognized dog breeds that provide the foundation for investigating the genetic basis of phenotypic diversity. Typically, breed standard phenotypes such as stature, fur length, and craniofacial structure are analyzed in genetic association studies. However, such analyses are limited to the assayed phenotypes, leaving difficult to measure phenotypic subtleties potentially overlooked. In this study, the genotype-first approach was adapted to the dog genome to investigate coding variation from over 2000 dogs, leading to discoveries of new mutations related to craniofacial morphology and stature. Breed-enriched variants were prioritized according to gene constraint, which was calculated using a mutation model derived from trinucleotide substitution probabilities in the dog. Among the discovered variants was a splice-acceptor mutation in *PDGFRA* associated with bifid nose, a characteristic trait of Çatalburun dogs, implicating the gene’s role in midline closure, and a frameshift mutation in *LCORL* associated with large canine body size, thus highlighting the importance of allelic heterogeneity in selection for breed traits. Most priority variants were not associated with genomic signatures for breed differentiation, as these regions were enriched for constrained genes intolerant to nonsynonymous variation, suggesting a model of breed phenotype diversification based on regulatory changes to essential genes. Identification of trait-associated variants in dogs informs new biological roles for genes. Improved collection of breed disease risk data, along with increased breed representation, will drive further discoveries.

## Introduction

The genomes of domestic species hold immense promise for the discovery of novel genotype-phenotype associations (Andersson and Purugganan 2022; Buckley and Ostrander 2024). The domestic dog comprises over 350 breeds (Rogers and Brace 1995; Fogle 2000), many of which exhibit extreme levels of phenotypic diversity, including those associated with morphology, behavior, and disease susceptibility (Ostrander et al. 2017; Leeb et al. 2023).

Identifying genetic variants that drive canine phenotypic variation will further define gene functions. Genetic association studies typically require an initial and robust characterization of a phenotype, a practice that ultimately limits the scope of potential findings to only those that can be recognized and sufficiently defined.

Alternatively, genetic associations can be identified using a “genotype-first” approach, where the first step is to identify candidate genotypes with potential phenotypic impacts (Stessman et al. 2014; Wilczewski et al. 2023). Based on the impacted genes, including predicted functions and characteristics of individuals with candidate variants, such as breed disease prevalence, researchers can generate new hypotheses for potential genotype- phenotype associations. In turn, this spurs additional collection of comprehensive phenotypes relevant to the trait of interest and allows for the hypothesized associations to be tested.

The genotype-first approach was initially formulated to investigate the genetic basis of phenotypic variation associated with complex human disorders which are frequently polygenic, non-coding, incompletely penetrant, and influenced by environmental factors (Stessman et al. 2014; Wilczewski et al. 2023). Complex traits often vary widely in presentation, and the presence of disease subtypes can require information regarding family history, disease presentation, treatment response, and histology. Genotype-first approaches, however, focus on classifying disease subtypes according to profiles of causative/pathogenic genetic variants. This alternate structure of disease classification can then guide comprehensive clinical phenotyping, making it possible to determine how specific alleles contribute to disease variability (Stessman et al. 2014; Leitsalu et al. 2021; Wenger et al. 2021; Park et al. 2022; Garcia-Pelaez et al. 2023; Wilczewski et al. 2023).

The adoption of the genotype-first approach in complex disease research has been largely driven by extensive genomic data collection in case cohorts, availability of electronic health records, and ability to recontact patients, supporting the identification of causative/pathogenic variants among cases and facilitating the collection of comprehensive phenotypes (Lyon et al. 2023; Safonov et al. 2023; Wilczewski et al. 2023; Wright et al. 2024). In canine genomics, there is far less availability of genomic and phenotypic resources. However, a rigid population structure, paired with extreme phenotypic diversity across breeds means that genotype-first approaches, when applied to dogs, can be used to ask different kinds of questions, specifically addressing the roles of specific genetic variants in shaping not only disease risk, but the striking differences in body morphology and behavior that characterize different breeds. By definition, dog breeds are closed populations, where members of a breed are only crossed with those of the same breed. Thus, every registered dog is descended from a small number of founders and shares extensive genetic homogeneity (Mellanby et al. 2013; Lampi et al. 2020). Breed members have multigeneration pedigrees and are judged against morphologic standards that further enforce genetic homogeneity, resulting in a reduction in the locus heterogeneity that typifies human traits, where dozens of genes may be foundational for a given complex phenotype. The ease of association studies in dogs is further enhanced by the fact that most breeds were developed within the last 200 years, such that there are minimal numbers of generations for meiotic recombination to occur (Worboys 2018). This unique population structure means that breed membership can be used as a surrogate when mapping breed-associated morphologic traits, thus avoiding the time and expense of collecting individual metrics (Gough et al. 2018; Plassais et al. 2019; Dutrow et al. 2022; Morrill et al. 2022).

In humans, candidate trait and disease variants are generally identified using a combination of strategies including multiple species alignment, protein structure predictions, gene mutation tolerance, and machine learning (Ng and Henikoff 2001; Adzhubei et al. 2010; Havrilla et al. 2019; Rentzsch et al. 2019; Karczewski et al. 2020; Cheng et al. 2023; Gao et al. 2023; Sullivan et al. 2023). Most such tools benefit from extensive catalogues of human variation featuring large amounts of clinical and personal data (Landrum et al. 2014; Lek et al. 2016; Bycroft et al. 2018; All of Us Research Program Genomics 2024). Although efforts to catalogue trait-associated genetic variation exist for domestic mammals (Nicholas et al. 1995; Plassais et al. 2019; Nicholas 2021; Meadows et al. 2023), the resources for extensive databases are limiting. While mapping functional gene/genomic annotations over from humans can theoretically be used for improving variant annotation, it is estimated that ∼13% of human pathogenic variants are likely benign in dogs (Gao et al. 2023). Systems such as the dog may, therefore, be rife with lost opportunities for identifying genes and variants that play fundamental roles in mammalian biology. Nevertheless, the recent Dog10K release of whole genome sequence (WGS) data from over 2000 dogs (Ostrander et al. 2019; Meadows et al. 2023), inclusive of 321 breeds and populations, means the time is ripe to use genotype-first methods, in combination with measures of functional and evolutionary constraint, to identify sequence level variants with a high probability of impacting gene function un the dog

This study focuses, specifically, on identifying genotypes that are rare, overall, in dogs but common within a single breed or groups of related breeds that share a particular trait. If the genotype highlights a variant of potential impact, it is selected for additional study. We characterize coding variation across 2,291 canine genomes from over 200 phylogenetically informed breeds/subpopulations. We use a mutation model to investigate the accuracy of three different gene/site constraint metrics for predicting variant phenotypic impacts. We then combine constraint metrics with variant data to apply the genotype-first approaches to canines, identifying several candidate trait-associated variants. Finally, we investigate the contribution of coding variation to breed differentiation, identifying the importance of constrained genes for shaping phenotypic differences between breeds.

## Results

### Canine genetic population structure and coding sequence constraint

To represent the breadth of canine coding variation, we combined the Dog10K cohort with a panel of dogs compiled from WGS data from the sequence read archive (SRA). After extensive sample and variant filtering, the combined dataset retained 821,202 coding sequence (CDS) sites with derived alleles from across 2,291 samples, including both splice donor and acceptor sites (**Fig. 1A**) (**Supplemental Table S1**) (**Methods**). Using the final dataset of 2,018 breed dogs, 219 village dogs, 53 wolves, and one coyote we further refined the dataset to 611,644 single nucleotide variants (SNVs), 28,970 deletions, and 28,123 insertions. Approximately 40% of all CDS SNVs occur only once in the dataset, indicating a vast landscape of uncharacterized rare variation within canids (**Fig. 1B**). We also identified 16,974 multiallelic sites, which consisted of 12,677 SNVs, representing 2.1% of all SNVs within the entire dataset, thus highlighting the extent of data loss from filtering for biallelic variants only, which can be expected to increase as datasets continue to grow (**Fig. 1C**). Dog breeds were also characterized as monophyletic clusters using a phylogenetic clustering approach (**Methods**) (**Supplemental Data S1**). A total of 200 breeds/groups with five or more WGS samples were identified (**Supplemental Table S2**).

**Figure 1:**
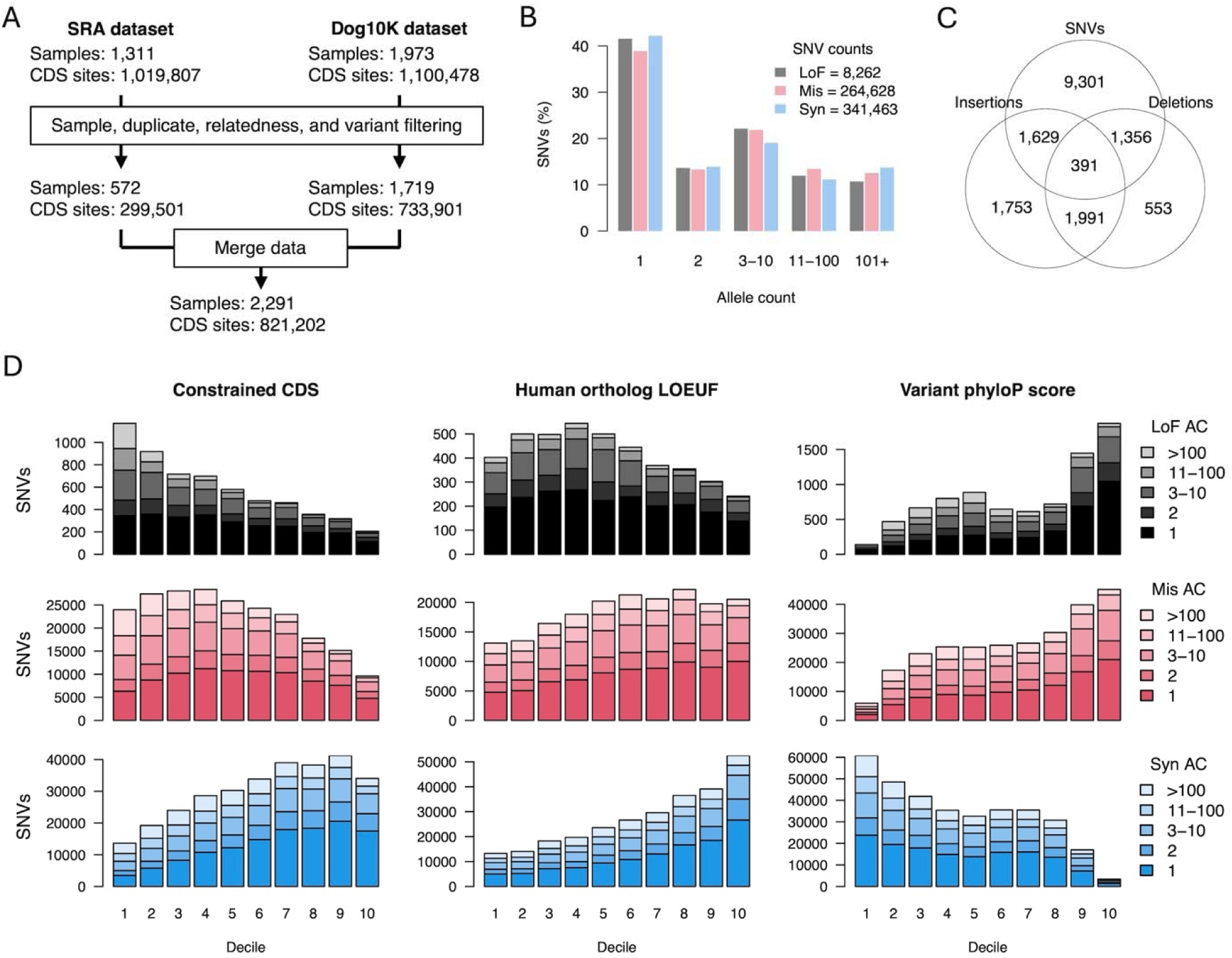
Coding variation and population genetics of 2,291 canine samples. A) Variant data from SRA and Dog10K were filtered and combined to create a single dataset used throughout the analysis. B) Allele count distribution of mutation types. C) Multiallelic variants occupying the same sites. D) The allele frequency distributions of SNVs according to mutation type and constraint level for three different constraint metrics. Constraint is expressed as a decile, where the highest deciles represent the most constrained genes/sites.

Variants were annotated according to three different constraint metrics: constrained CDS, human ortholog LOEUF (loss of function observed over expected upper fraction), and variant phyloP score. Constrained CDS and variant phyloP score both represent evolutionary conservation derived from the Zoonomia consortium’s alignment of 240 mammals. Variant phyloP score is the level of conservation of an individual nucleotide position. Constrained CDS is the fraction of nucleotides within a gene’s CDS that have a phyloP score >2.27 (genome-wide FDR > 0.05) (Christmas et al. 2023; Sullivan et al. 2023). Human ortholog LOEUF indicates whether a human-dog ortholog is depleted of predicted loss of function (LoF) variation in humans (Karczewski et al. 2020), where LoF here specifically represents nonsense, splice disruptor, and frameshift variant annotations (MacArthur et al. 2012). The values used are from GNOMAD v4.1.0 and are based on expected rates of LoF variation from over 800,000 humans (Chen et al. 2024). The key differences between the constraint metrics are 1) whether they reflect evolutionary constraint (constrained CDS and variant phyloP score) or functional variation constraint (human ortholog LOEUF), and 2) whether the metric represents an entire gene (constrained CDS and human ortholog LOEUF) or a specific base pair (variant phyloP score). These differences are important, as evolutionary-based metrics, which depended on across species sequence conservation, are blind to DNA sequences with essential lineage- specific biological roles, and gene-based metrics have too low resolution for evaluating selection effects on gene regions or protein domains.

To characterize each constraint metric in dogs, we analyzed the distribution of SNV mutation types and allele counts according to constraint level (**Fig. 1D**). As expected, genes under the greatest constraint, according to constrained CDS and human ortholog LOEUF, had the lowest fraction of LoF variants. Conversely, LoF variants were most common among sites with high variant phyloP scores, the opposite of what was observed for gene-based metrics.

This outcome is caused by codon position effects on phyloP scores. Positions susceptible to LoF substitutions are going to be the most conserved across species, indicating that LoF variants are going to occur more frequently at these positions (Pollard et al. 2010; Sullivan et al. 2023). The inverse of these effects is observed with synonymous variants, which are most abundant in highly constrained genes and almost completely absent from highly constrained sites, due to their often-negligible impact on fitness.

In dogs, allele count distributions, a proxy for allele frequencies, also varied according to constraint level. LoF SNVs in constrained genes/sites were usually rare, while common LoF SNVs (AC >100) were more frequently found within genes/sites not under strong constraint. Although low allele frequency and density of functional variants is an indicator of negative selection acting on deleterious mutations, the observed difference in SNV density across constraint metrics suggests that the utility of each metric for variant prioritization may vary significantly.

### A framework for evaluating gene sequence constraint metrics

Genomic sequence constraint metrics are often used to predict whether specific mutations are likely to have deleterious impacts. Characteristic breed traits in dogs are often phenotypic extremes and can be linked to deleterious health outcomes, such as in brachycephalic breeds (Marchant et al. 2019; O’Neill et al. 2020). Sequence constraint, therefore, has potential for identifying trait-causing variants in dogs. However, the constraint metrics used here do not measure canine coding constraint directly. Instead, evolutionary-based metrics identify sites and genes conserved across many species and are blind to lineage- specific effects. Similarly, LOEUF values were calculated in human genes, and are not only blind to canid lineage-specific selection but also capture human lineage-specific effects that may not be relevant to canines. We therefore evaluated the capacity of each constraint measure to predict gene tolerance to mutations within dogs by measuring their concordance with the ratio of observed to expected mutation rates in canine genes.

Calculation of the observed mutation rates for this analysis must only capture negative selection effects and no other population genetic effects. One approach is to use recent spontaneous mutations, whose presence in the genome is unaffected by population dynamics. Similarly, expected mutation rates must also be unbiased. The observed genetic variation within dogs is a result of their complex demographic history, including recurrent population bottlenecks and strong selective sweeps (Marsden et al. 2016). It is difficult to determine whether a LoF mutation has persisted because it has a low deleterious impact or because it has a high deleterious impact but is in linkage disequilibrium (LD) for a desired trait. Canine trinucleotide substitution rates and coding sequence composition was therefore used to build a canine gene mutation model, an approach similar to calculating LOEUF expected mutation rates (Samocha et al. 2014) (**Methods**).

Singleton variants represent the best candidates for recent spontaneous mutations. To further enrich for recent mutations, dogs with highly represented ancestry were identified using individual singleton counts, as excess singletons per individual is an indicator of underrepresented ancestry. Most dogs, ∼70%, carried <40 singletons each within their coding sequences, allowing this value to serve as a cutoff (**Supplemental Fig. S1A**). These dogs belonged to breeds that had significantly more members in the dataset (greater Wilcoxon rank sum test; P-value < 10^-15^), and larger fractions of their genomes were within runs of homozygosity (F_ROH_) (greater Wilcoxon rank sum test; P-value < 10^-15^) than was observed for individuals with >40 singletons (**Supplemental Fig. S1B**). Only 11.2% of all singletons were inferred to be recent spontaneous mutations by this method, and most low-frequency variants were contributed by a minority of individuals with poorly represented ancestry (**Supplemental Fig. S1C**).

Next, the canine mutation model was evaluated by testing whether it could predict the number of synonymous mutations per gene. Since synonymous mutations typically have little biological impact on coding sequence, their observed frequency should be highly correlated with gene mutation probabilities. The Pearson correlation coefficient was used to evaluate model fit. A total of 8,660 autosomal synonymous singletons were extracted from dogs with highly represented ancestry and used as observations. A modest correlation of 0.53 was achieved, yet due to the low number of observations, there was likely an insufficient number of events for expected trends in mutation accumulation to emerge.

Sparsity in observations was addressed first by relaxing criteria for defining recent spontaneous mutations, which involved increasing maximum allele count thresholds and including variants from all dogs, and second by grouping genes based on mutation probabilities. The 95% confidence intervals for expected fit were also calculated, allowing the mutation model to be evaluated as a departure from expected fit values, which is less sensitive to the number of observed events than the Pearson correlation coefficient. Although relaxing criteria for defining recent mutations led to increased correlations, the departure from expected model fit also grew (**Fig. 2A**), indicating that inclusion of additional variants to the analysis diluted the fraction of true recent spontaneous mutations. The alternative strategy of placing genes into groups based on mutation probability was more successful for testing model fit. For 1000 groups or less, correlations greater than 0.9 were reached and the model fit remained within expected ranges, unlike a competing model based on CDS length (**Fig. 2B**).

**Figure 2:**
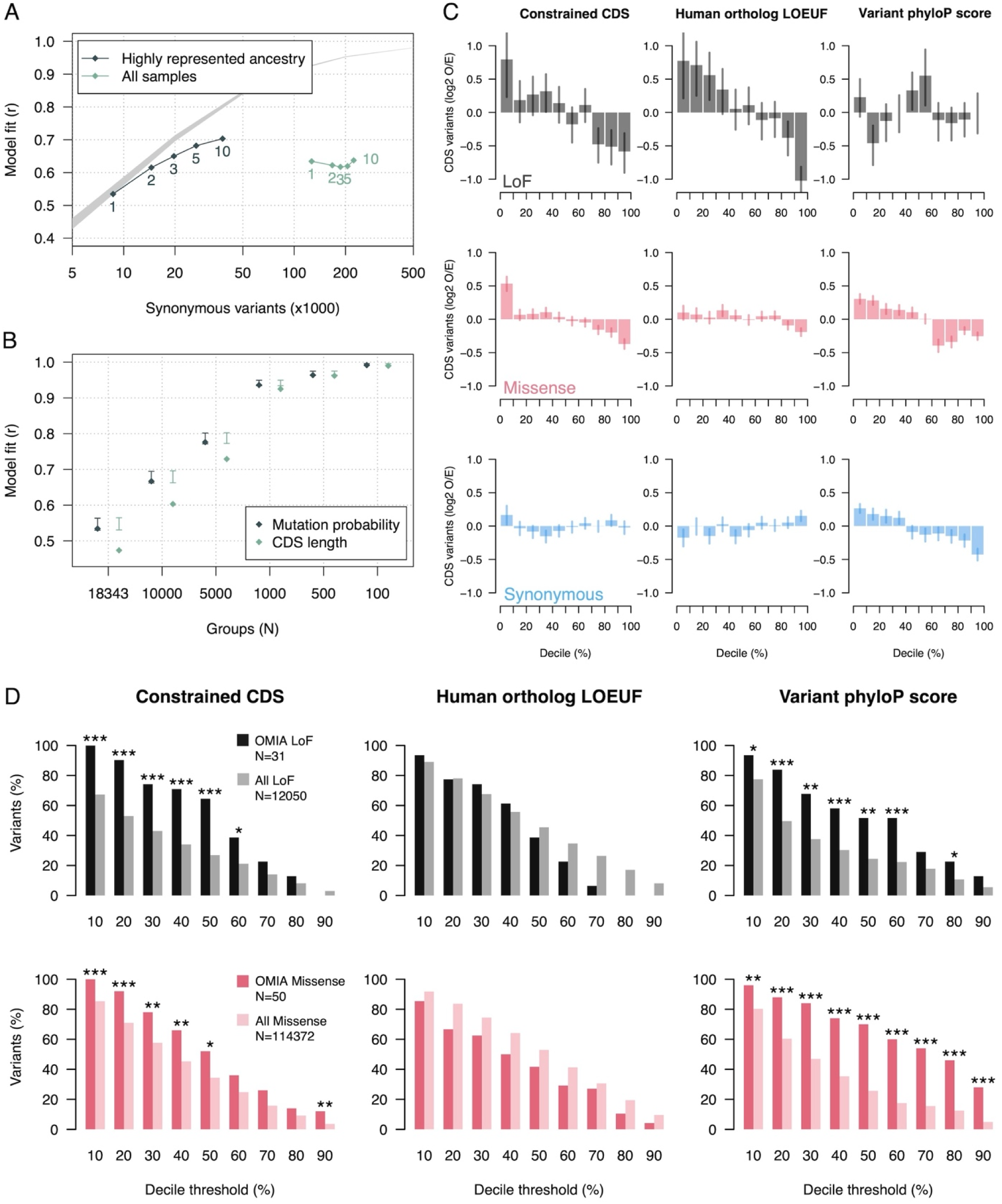
Recent spontaneous mutations in dogs and selection effects on constrained regions. A) Pearson correlation coefficient between observed rare variants and gene mutation probabilities. Variants are organized by the group of samples they were observed in and the maximum allele count threshold for defining rare variants, which is displayed near each datapoint. The grey band represents the 95% expected interval of the model fit based on simulated accumulation of synonymous mutations. B) Pearson correlation coefficient of rare variant frequency and gene mutation probability across gene groups. Genes were grouped according to their mutation probabilities. The x-axis shows the number of groups used to calculate correlation coefficients. The number of groups decreases as group sizes get larger. Two models are tested, one is the mutation probabilities calculated using the mutation model and the other is mutation probabilities according to gene CDS length. The 95% confidence interval for the expected model fit are also displayed. C) Mutation frequency in genes ranked by different constraint metrics. The expected number of mutations is calculated following the mutation model. The observed number of mutations are from singletons in dogs with highly represented ancestry. Error bars indicate whether the observed mutation frequency is outside the 95% expected bounds. D) Enrichment of OMIA variants after variant/gene constraint filtering. Vertical bars show the fraction of OMIA and all other variants that remain after implementing each filtering threshold. Significance was determined using a Fisher test. “*” indicates P-value < 0.05, “**” indicates P-value < 0.01, and “***” indicates P-value < 0.001.

Together, these analyses show that accumulation of recent spontaneous synonymous mutations across genes closely follows expected mutation probabilities, demonstrating that it is possible to identify mutations in dogs where presence in the genome is largely independent of canine population history, and that the proposed canine mutation model can accurately calculate expected mutation frequencies for a given number of observations. This framework can therefore be used to determine if genes have significantly fewer nonsynonymous mutations than expected, thus detecting negative selection effects in the dog.

### The functional constraint of dog genes

Using 907 LoF and 14,947 missense singleton mutations from dogs with highly represented ancestry, we tested whether the constraint metrics were associated with negative selection effects. For gene-based metrics, dog genes were ranked and grouped into deciles by constraint level, and mutation probabilities were summed accordingly. For variant phyloP scores, a site-based metric, the individual CDS positions, were grouped into constraint deciles, and base-pair mutation probabilities were summed according to each decile. Also, since phyloP scores are heavily stratified by mutation type (**Fig. 1D**), breaks between phyloP score deciles were made mutation type specific, thus balancing the expected frequencies of mutations across the full spectrum of phyloP constraint.

Results show that gene-based constraint metrics in dogs are significantly associated with intolerance to LoF mutations (**Fig. 2C**). The top three deciles for constrained CDS and top two deciles for human ortholog LOEUF were significantly depleted of LoF mutations, indicating that for this group of genes a significant portion of the expected LoF mutations likely had deleterious impacts on fitness. The strongest selection effects were observed in the top human ortholog LOEUF decile, where only half the expected number of LoF mutations were observed (**Fig. 2C**). Conversely, no significant negative selection effects were observed for LoF mutations among the top phyloP constraint deciles, indicating that evolutionary conservation across species does not always correlate with mutation impacts on fitness.

In dogs, gene level constraint was also associated with intolerance to missense mutations (**Fig. 2C**). These effects were less pronounced than LoF selection effects, as functional consequences from missense mutations are often less severe. For variant phyloP scores, intolerance to missense mutations was observed across the top four constraint deciles. For synonymous mutations, no significant negative selection effects were observed amongst the top gene-based constraint deciles, consistent with synonymous mutations having mostly negligible impacts on gene function. Unexpectedly, top phyloP score deciles appeared significantly intolerant to synonymous substitutions. However, since phyloP scores are site- based, the observed intolerance may be in response to synonymous mutations that can cause cryptic-splice sites, impact mRNA stability, have reduced tRNA availability, or induce change in functionally relevant GC content (Sarkar et al. 2022; Gudkov et al. 2024).

We next tested whether constraint metrics were associated with known canine trait variants. We increased the minimum threshold for defining constraint by decile increments and measured enrichment of trait-associations within the remaining fraction variants. Canine trait- associated variants were extracted from the online mendelian inheritance in animals (OMIA) database (**Supplemental Table S3**), a catalogue of inherited disorders and other traits, and associated genes in non-model animal species (Nicholas 2021; Nicholas 2024) (https://www.omia.org/home/). Although OMIA is the largest collection of animal trait-associated variants, supporting evidence for trait-associations vary widely, ranging from candidate alleles found in single individuals, to mutations shown to be causative in functional studies. We focused only on OMIA variants with an allele count > 2 within the combined cohort. This cutoff was used to ensure that we would only analyze variants that could be identified using genotype-first approaches. Altogether, 31 out of 305 LoF and 50 out of 154 missense OMIA variants met these criteria (**Supplemental Table S3**).

For evolutionary-based constraint metrics, we detected enrichment of both LoF and missense OMIA variants (**Fig. 2D**). However, in most cases increases to the minimum constraint threshold beyond a certain point led to a loss of OMIA enrichment (**Supplemental Table S4)**. Similarly, no enrichment of LoF OMIA variants were detected for human ortholog LOEUF constraint, which was associated with the strongest negative selection effects. This outcome indicates that OMIA LoF variants likely do not have severe deleterious impacts on fitness. Such variants would be unlikely to persist in canine populations, unlike the variants analyzed here, which were all present in multiple dogs within the cohort.

### Genotype-first approaches applied to canine variation

For most dogs in this dataset, the only individual information available was sex and breed. We therefore focused on genotypes that were enriched in a particular breed. We also investigated the breed specificity of OMIA variants across a range of minimum breed frequency cutoffs. We found that increases to the minimum breed allele frequency cutoff result in fewer discoverable OMIA variants. At a 20% minimum breed allele frequency, the fraction of the remaining variants found in only one breed is at its highest (**Supplemental Fig. S2**). For the remainder of the analyses, we therefore only considered variants with a minimum breed allele frequency >20% in at least one breed.

Lists of potential LoF and missense candidates for canine traits were defined using each constraint metric. Where possible, cutoffs were selected at the points where negative selection effects could be detected (**Fig 2C**). When there was no such effect, such as for variant phyloP scores and LoF mutations, the highest percentile cutoffs where OMIA enrichment was observed were used instead (**Fig 2D**). Variant lists were filtered using a minimum breed allele frequency of 20%, a minimum allele count of three, a minimum breed allele number of six, and a maximum non-breed allele frequency of 5%. Variant lists were then ranked according to the number of breeds the variant was found in and the proportion of individuals in those breeds carrying the variant (**Fig 3**.) (**Supplemental Table S5-S10**).

**Fig 3:**
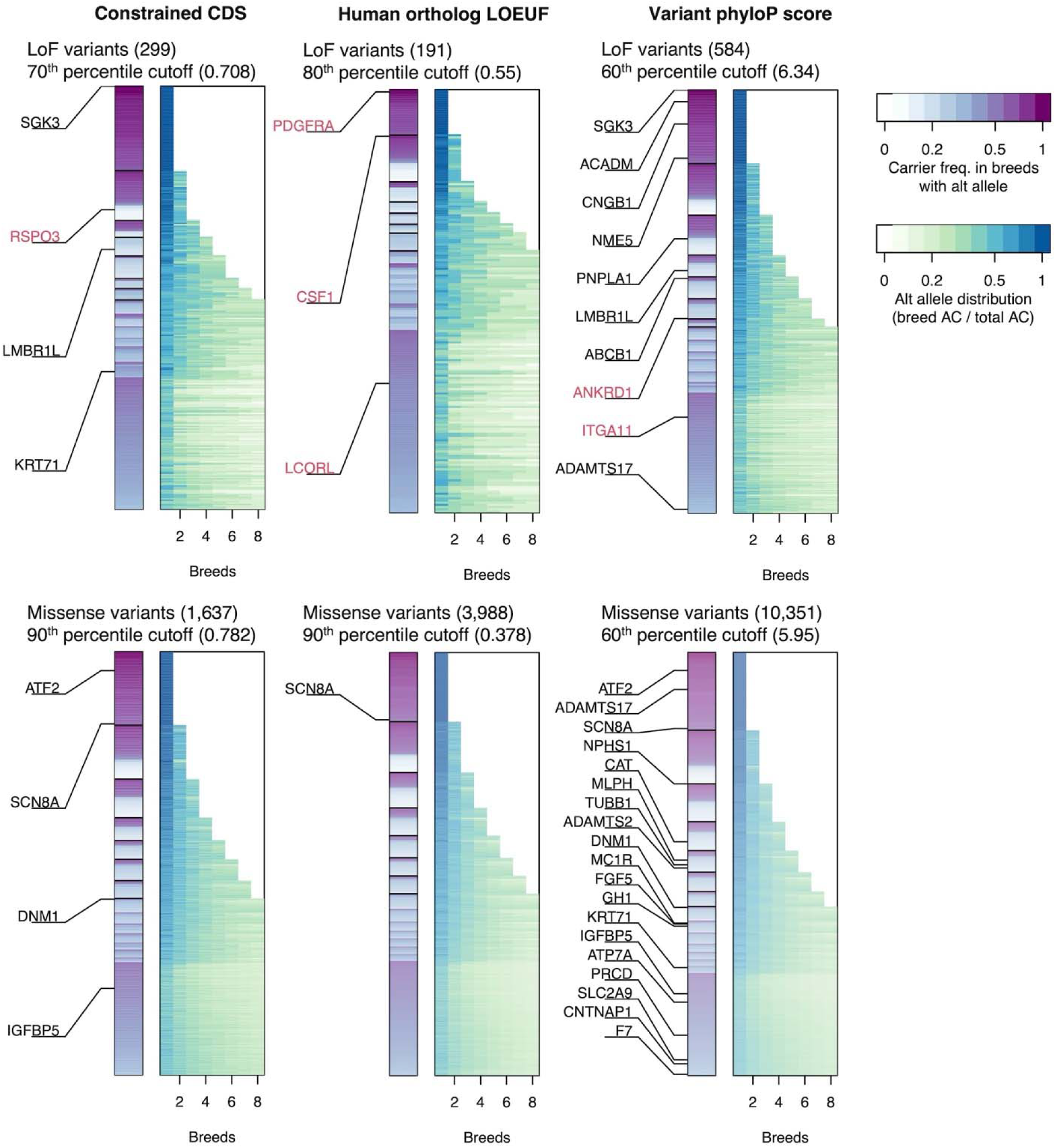
List of candidate LoF/missense variants for trait associations. Variants are ordered along the y-axis according to number of breeds they are found in and the proportion of individuals in those breeds carrying the variant. Breeds carrying the variant are ordered along the x-axis according to the proportion of the alt alleles they are carrying. Gene names printed in black text contain OMIA variants. Gene names in red are non-OMIA variants that were identified after manual curation of the variant list. Variant type and constraint cutoff are displayed for each panel.

For LoF constraint, the number of variants returned after filtering by breed allele frequency ranged from 191 for human ortholog LOEUF to 584 for variant phyloP score. For missense mutations, the number of returned variants ranged from 1,637 for constrained CDS to 10,351 for phyloP score (**Fig. 3**). Since LoF constraint filtering returned fewer variants, and is more highly associated with negative selection effects, the associated variant lists were manually investigated to identify new candidate genotype-phenotype relationships. Based on their potential for phenotypic impact, a total of six variants were selected for comment (**Table 1**).

**Table 1:** Breed genotypes and allele frequencies of select candidate variants.

| Variant | Breed | AF | Ref | Het | Hom |
| --- | --- | --- | --- | --- | --- |
| chr13:47440160, A/C, *PDGFRA*, splice acceptor | Catalburun | 0.5 | 0 | 3 | 0 |
| chr1:66494619, C/T, *RSPO3*, stop gained | Irish Water Spaniel | 0.33 | 2 | 4 | 0 |
|  | Unassigned | 0 | 324 | 1 | 0 |
| chr6:42492671, C/A, *CSF1*, stop gained | Chow Chow | 1 | 0 | 0 | 6 |
|  | Elo | 0.1 | 4 | 1 | 0 |
| chr3:91871030, AACAG/A, *LCORL*, frameshift variant<sup>1</sup> | Spanish Mastiff | 1 | 0 | 0 | 6 |
|  | Berger Picard | 0.77 | 0 | 5 | 6 |
|  | Briard | 0.67 | 0 | 4 | 2 |
|  | Petit Bleu de Gascogne | 0.58 | 1 | 3 | 2 |
|  | Bluetick Coonhound | 0.5 | 0 | 2 | 0 |
|  | Bracco Italiano | 0.5 | 3 | 0 | 3 |
|  | Spinone Italiano | 0.43 | 2 | 4 | 1 |
|  | Pyrenean Mastiff | 0.36 | 3 | 3 | 1 |
|  | Leonberger | 0.32 | 9 | 8 | 2 |
|  | Beauceron | 0.3 | 2 | 3 | 0 |
|  | Mastiff | 0.28 | 5 | 3 | 1 |
|  | German Hound | 0.25 | 3 | 3 | 0 |
|  | Griffon Nivernais | 0.25 | 2 | 2 | 0 |
|  | Ariege Pointer | 0.2 | 3 | 2 | 0 |
|  | Karakachan | 0.2 | 3 | 2 | 0 |
|  | Picardy Spaniel | 0.2 | 3 | 2 | 0 |
| chr30:32785671, ACT/A, *ITGA11*, frameshift variant<sup>1</sup> | Pug | 1 | 0 | 0 | 3 |
|  | Japanese Chin | 1 | 0 | 0 | 4 |
|  | Pekingese | 1 | 0 | 0 | 4 |
|  | Shih Tzu | 1 | 0 | 0 | 4 |
|  | Tibetan Spaniel | 1 | 0 | 0 | 4 |
|  | Boston Terrier | 1 | 0 | 0 | 6 |
|  | French Bulldog | 1 | 0 | 0 | 6 |
|  | Lhasa Apso | 1 | 0 | 0 | 6 |
|  | Chihuahua | 0.36 | 4 | 1 | 2 |
|  | Biewer Terrier | 0.3 | 2 | 3 | 0 |
|  | Portuguese Podengo Pequeno | 0.25 | 1 | 1 | 0 |
|  | Russian Tsvetnaya Bolonka | 0.25 | 3 | 3 | 0 |
|  | Chesapeake Bay Retriever | 0.21 | 4 | 3 | 0 |

<sup>1</sup>Only breeds with AF $\geq 0.2$ are displayed

The most interesting candidate was a splice-acceptor within the *PDGFRA* gene that was found in a heterozygous state in all three Çatalburun dogs within the cohort. Members of this breed, also called the Turkish pointer, are notably characterized by a split/bifid nose (**Fig 4A**) (Yilmaz et al. 2012). Indeed, the English translation of “Çatalburun“ is “forked nose” (Özarslan et al. 2021). Mouse knockout models demonstrate that *PDGFRA* plays an important role in medial nasal process development and palatogenesis (He and Soriano 2013; Qian et al. 2017), thus aligning with the hypothesis that the *PDGFRA* splice-acceptor variant causes the Çatalburun’s bifid nose phenotype. The variant was genotyped in fourteen additional Çatalburun dogs, although phenotypes were not available, the variant was heterozygous in all but one dog which was homozygous for the reference allele (**Supplemental Table S11**).

**Figure 4:**
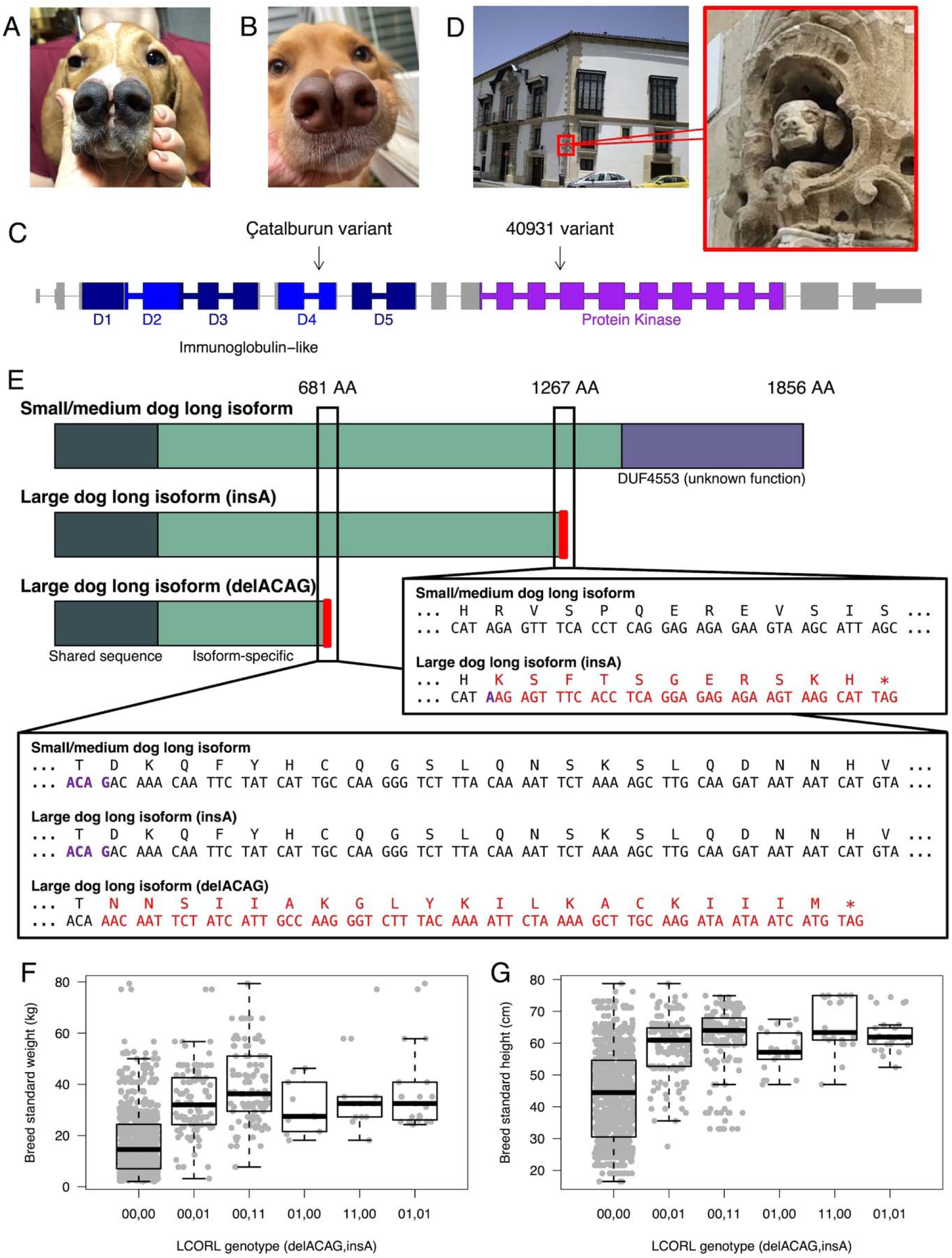
Phenotypic variant discovery. A) Çatalburun dog with distinctive bifid nose phenotype. B) Golden retriever mix (Dog ID: 40931) displaying a bifid nose phenotype and is heterozygous for an additional *PDGFRA* splice-acceptor variant. C) Canine *PDGFRA* transcript showing the locations of each splice acceptor variant found within dogs with orofacial clefts. The first variant is the çatalburun variant with the second variant belonging to dog 40931. D) Photograph of dog statue with split-nose phenotype. The statue is located at the Palace of Bertemati in Andalusia, Spain. The palace was constructed in the 18^th^ Century. E) Mutations found in the long isoform of *LCORL*. Early termination of translation and loss of the DUF4553 domain caused by frameshift mutations appears to be associated with larger dog size (> 25kg/>55cm). F) *LCORL* genotypes and breed standard weight. G) *LCORL* genotypes and breed standard height.

To further investigate the genetics of this phenotype, we recruited and performed WGS on eight additional bifid nose dogs, all lacking any apparent Çatalburun ancestry. Six of the dogs were unrelated to one another, while two were litter mates. An unaffected parent of the two affected siblings was also in included (**Supplemental Fig. S3**). Variants were prioritized according to evolutionary constraint and potential functional impact on orofacial cleft associated genes. Twelve heterozygous candidate variants were identified across all eight non-Çatalburun bifid nose dogs (**Supplemental Table S12**). One sample, 40931 (**Fig. 4B**), carried a heterozygous insertion that also disrupted a *PDGFRA* splice-acceptor but at a different position (**Fig. 4C**). When considering all 2,291 dogs in the entire cohort, only those with the bifid nose phenotype carried LoF mutations in *PDGFRA*, thus defining the existence of a naturally occurring model for study of *PDGFRA’s* role in craniofacial development and midline closure. An additional breed, the Pachón Navarro, which originates in Spain, is also known for its bifid nose phenotype. Unfortunately, no samples from this breed were available for study. Interestingly, however, masonry on the outside of the 18^th^ century Palace of Bertemati (Andalusia, Spain) shows a gargoyle-like dog with a bifid nose (Cobo 2017), attesting to the continuous selection for this trait in the Pachón Navarro breed (**Fig. 4D**).

Another variant with potential implications on craniofacial morphology is a frameshift within *ITGA11* (**Table 1**). It is fixed in small brachycephalic breeds, and mouse knockouts of the gene demonstrate its association with small size and craniofacial abnormalities, specifically the eruption of incisors (Popova et al. 2007). In dogs, an association between the locus and brachycephaly was described previously but was sensitive to corrections for canine size (Schoenebeck et al. 2012; Marchant et al. 2017). The variant could cause a range of complex phenotypes, contributing to both size and craniofacial structure. However, additional functional analyses are needed.

A new mutation within *LCORL*, a previously described canine size-associated gene (Hayward et al. 2016; Plassais et al. 2019), was also identified. The previously described variant, c.3768_2769insA, causes a frameshift and premature stop codon in the long isoform of the gene. The variant is one of the largest contributors to canine size, explaining ∼15% of the variance in canine standard breed weight (Plassais et al. 2019). Here, a newly identified *LCORL* variant, c.1978_1981delACAG, a deletion upstream of the previously described insertion, also causes a frameshift and premature stop (**Fig. 4E**). The c.1978_1981delACAG variant was fixed in Spanish Mastiffs, an extremely large breed whose standard has no upper limit on size, and segregates in several other large breeds, i.e., those weighing > 25kg (**Table 1**). At an allele frequency of 2.9%, the newly identified c.1978_1981delACAG variant is less common across the entire cohort than the c.3768_2769insA, which has an allele frequency of 31.2%. Among large dogs, the frequency of the c.1978_1981delACAG is 6.4% and 37.9% for c.3768_2769insA. Both alleles were also significantly associated with both breed standard weight (**Fig. 4F**) and height (**Fig. 4G**) (P-value < 10^-10^). There were 30 instances where dogs had one copy of each allele and no instances of a dog having two copies of one allele and one or two copies of the other, indicating that the c.3768_2769insA and c.1978_1981delACAG alleles are most likely on different haplotypes.

Another variant that may correspond to a breed trait is a stop-gain at position p.Glu391* in *colony stimulating factor 1* (*CSF1*). The variant was homozygous in all six unrelated Chow Chows (Chow), heterozygous in one of five Elos, and absent from two Eurasiers, the latter two breeds being the product of Chow admixture (**Table 1**), thus indicating a high degree of breed specificity. The Chow is a medium-sized spitz-type breed that traces its origins to China (Yang et al. 2017). It is characterized by its thick double coat, black pigmented tongue, straight hind legs, broad square skull and muzzle, and supernumerary third incisors (Yang et al. 2017). The region containing *CSF1* is under strong positive selection in Chows (Yang et al. 2017), suggesting genes in this region contribute to the breed’s characteristic phenotypes. The p.Glu391* mutation is located in exon six of the secreted isoform of the CSF1 protein (Cosman et al. 1988). Rodent knockout models implicate *CSF1* in skeletal and craniofacial development, specifically a domed skull, missing incisors, short thickened limb bones, fertility, and macrophage differentiation (Marks and Lane 1976; Yoshida et al. 1990; Dai et al. 2002; Van Wesenbeeck et al. 2002; Sehgal et al. 2021). While the precise function of the p.Glu391* mutation in dogs is unknown, the rodent phenotypes of a domed skull and altered number of incisors, suggest p.Glu391* may contribute to the Chow’s broad square skull and two additional incisors, which contrast sharply with the Eurasier and Elo breed requirements.

One variant with potential medical consequences was a stop gain within *RSPO3* and was found in a heterozygous state in four out of six Irish Water Spaniels and one Xoloitzcuintle, the latter being a hairless Mexican breed whose members were largely unassigned to a breed/group during phylogenetic clustering (**Table 1**). This mutation is of particular interest as a 167 bp insertion in the 3’UTR of a gene from the same family, *RSPO2*, is responsible for “furnishings”, defined as the long moustache and increased eyebrow hair observed in breeds like the schnauzer (Cadieu et al. 2009). Depletion of Rspo3 within the mouse dermal papillae causes delayed hair regrowth, indicating it is required to activate proliferation of dermal stem cells within the hair follicle, although *RSPO2* may provide some functional redundancy (Hagner et al. 2020). Since the *RSPO3* mutation in Irish water spaniels is not found within every breed member, it is unlikely to be connected to a breed-characteristic trait. Rather, the variant may explain why some Irish Water Spaniels suffer alopecia from the trunk or thighs as they age, which occurs with high frequency (Cerundolo et al. 1999).

### Coding variation within genomic regions exhibiting differentiation across breeds

Artificial selection for characteristic breed traits has left its mark on the dog genome. Within breeds, there are extended regions of homozygosity that are also associated with strong signatures of population differentiation between breeds (Akey et al. 2010; Vaysse et al. 2011; Parker et al. 2017a; Morrill et al. 2022). Moreover, trait-associated variants for breed characteristic traits, such as ear shape, body size, and sport-hunting ability are often within breed-differentiated genomic regions, where genetic variation is extremely stratified by breed (Boyko et al. 2010; Kim et al. 2018; Plassais et al. 2019). Like genotype-first approaches, breed- differentiated regions provide a genomic signature potentially associated with canine breed traits. However, unlike genotype-first approaches, breed-differentiated loci often encapsulate multiple genes, making it difficult to identify specific variants that contribute to breed characteristic traits. To enhance detection of trait-associated variants, potentially functional variants in constrained genes were analyzed for their association with signatures of breed differentiation.

Pairwise F_ST_ between 200 breeds/groups was calculated for 50kb genomic windows. Windows with F_ST_ > 0.8 signified extreme differentiation between a breed/group pair. Loci with > 100 extremely differentiated breed/groups pairs were classified as breed differentiated regions.

These regions were further refined using peak detection software (Locher 2024). Genomic segments of extended breed homozygosity were also identified and were also used to further refine breed-differentiated regions by providing an additional marker of artificial selection (**Methods**) (**Fig. 5A**). In total, there were 1,743 breed-differentiated regions, containing 2,085 genes (**Supplemental Table S13**). However, 92.3% and 55.2% of breed-differentiated regions were within 1 Mb and 100 kb of each other, respectively, indicating that most breed- differentiated regions are likely part of larger selective sweeps. Seventeen of 25 autosomal trait- associated loci described in Plassais et al. (2019) were located within 100kb of a breed- differentiated region, indicating that the regions identified here are associated with breed traits under selection (**Fig. 5B**). Population branch statistics from an analysis of 101 different breeds inclusive of >12 dogs were also used to identify breed-differentiated regions, capturing a total of 3,016 genes (Morrill et al. 2022). Of these, 726 were within breed-differentiated regions identified in this analysis.

**Fig 5:**
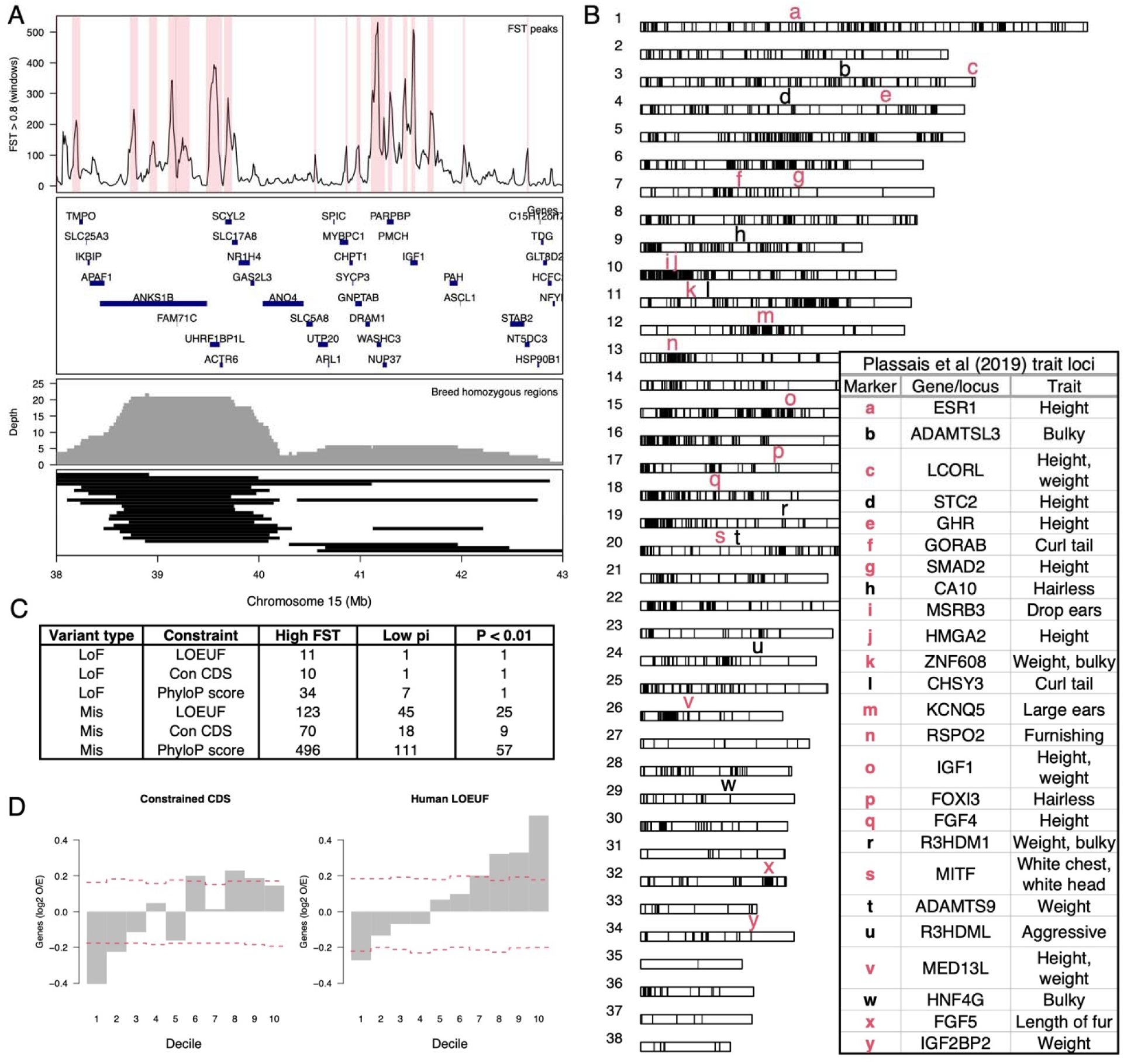
Non-coding variation at constrained genes drives breed differentiation. A) Identification of breed differentiated regions at chromosome 15 IGF1 locus. Genomic windows with F_ST_ > 0.8 between breed pairs were plotted across the genome, and peaks were then identified as regions of breed differentiation. High F_ST_ peaks are shown in the top panel and highlighted in red. The second panel shows genes found in the region. The third panel shows the density of regions of breed homozygosity that were identified as 1 Mb genomic regions with breed nucleotide diversity < 0.02. The final panel shows black bars as individual breed homozygosity stretches. B) Genome-wide display of breed-differentiated regions that overlap previously characterized trait loci from Plassais et al (2019), which are displayed as lower-case characters. Loci in red are within 100kb of a breed-differentiated region. C) Number of genes in highly differentiated regions carrying potentially functional variants that pass minimum constraint thresholds. The High F_ST_ column is the number of genes that carry variants of interest. The Low pi column is the count of genes that carry variants that are in breeds with low nucleotide diversity across the differentiated region. The final column is the number of genes with variants enriched in breeds that share the same haplotype within the differentiated region. D) Observed over expected frequency of genes at different constraint levels within highly differentiated regions.

There were between 10 and 34 genes across all breed-differentiated regions that contained LoF mutations with potential functional consequences (**Fig. 5C**). Of these, only two were found in breeds that exhibited signatures of differentiation at these loci, specifically the *CSF1* premature stop codon found in Chows (Table 1) and the *SGK3* mutation causing hairlessness in American Hairless Terriers (Parker et al. 2017b). Although not significantly associated with breed differentiation, a subset of mutations was within breed-differentiated regions and segregating in the dogs whose breeds contained long stretches of homozygosity. One example is a frameshift mutation caused by a single base pair insertion in *CYP4A11*, a monooxygenase involved in drug metabolism and synthesis of cholesterol, steroids and other lipids, associated with cardiovascular and cerebrovascular disease and hypertension in humans (Sugimoto et al. 2008; Yu et al. 2018) (**Supplemental Table S14**). The canine variant may also be associated with health outcomes in dogs and may persist because it is in LD with a trait under positive selection.

Altogether, 70 genes contained missense mutations with potential functional impacts that were significantly associated with breed differentiation (**Fig. 5C**). The strongest result was for a missense variant in *RYR1* (**Supplemental Table S14**), a calcium ion channel primarily expressed in the sarcoplasmic reticulum of skeletal muscle and involved with muscle contraction (Hernandez-Ochoa et al. 2015). The variant was almost exclusively found within Bull Terriers and American Staffordshire Terriers, two breeds with shared ancestry that are known for their musculature and agility. Another member of the gene family, *RYR3*, which is also involved with muscle contraction, is under positive selection in sport-hunting breeds (Kim et al. 2018).

The vast majority of breed-differentiated regions were not associated with nonsynonymous variants, indicating that breed selection may instead be acting upon non-coding variants that likely impact gene expression, which is consistent with most mapping studies for breed-defining traits and breed/lineage differentiation (Dutrow et al. 2022; Morrill et al. 2022). The genotype-first approach described here focuses on identifying variants with potential phenotypic impacts, prioritizing variants in constrained genes. Since breed- differentiated regions often represent loci under selection for breed characteristic phenotypes (Plassais et al. 2019; Morrill et al. 2022), these regions should also be comprised of constrained genes. We tested for enrichment across constraint deciles for gene-based metrics and found that human ortholog LOEUF constraint was significantly overrepresented (**Fig 5D**). This supports a model of phenotypic breed differentiation driven by regulatory variation of highly constrained genes rather than alterations to gene function from nonsynonymous mutations.

## Discussion

Artificial selection for desired traits has made the domestic dog one of the most phenotypically diverse mammals in the world (Wayne and vonHoldt 2012). Rigid population structure, enforced by breed standards, has promoted high levels of homozygosity within breeds while also maintaining isolation between breeds (Parker et al. 2004; Karlsson and Lindblad-Toh 2008). Together, these factors make the domestic dog an invaluable system for identifying genes related to breed characteristics, which are often morphological extremes (Ostrander et al. 2017; Buckley and Ostrander 2024). We therefore investigated the genetic basis of phenotypic breed differences using a genotype-first approach. Specifically, we used metrics for genomic constraint to identify variants with potential phenotypic impacts. We then ranked variants according to breed enrichment. Variants common to one or a few breeds but rare throughout the population are potential candidates for breed traits. This work provides significant advances to the field of canine genetics by: 1) defining a variant set whose presence in the genome is unaffected by canine population dynamics, thus making it possible to determine genes under negative selection; 2) evaluating three constraint metrics for prioritizing variants for potential phenotypic impacts; 3) identifying new CDS variants with strong impacts on craniofacial morphology and body size; and 4) applying genomic constraint to support a model of breed differentiation based on non-coding variation.

Using the genotype-first approach, we identified strong candidates for phenotypic impacts as previously unreported LoF variants in *PDGFRA* and *LCORL,* genes both under strong human ortholog LOEUF constraint. Two *PDGFRA* variants were identified, one was in Çatalburun dogs, a breed largely characterized by a bifid nose phenotype (Fig. 4A), and the other was in a Golden Retriever mix. This gene may therefore be one of a few for which functional mutations affecting midline closure occur, but are tolerated, as long as they are in the heterozygous state. In humans, a missense variant in this gene within a single family is associated with left non-syndromic cleft-lip and palate but is not fully penetrant (Yu et al. 2022). *PDGFRA* is a type III receptor tyrosine kinase that consists of an extracellular region with five immunoglobulin-like domains for ligand binding, a transmembrane helix, and an intracellular tyrosine kinase domain (Williams 1989; Chen et al. 2013). The Çatalburun variant occurs in the seventh intron of the gene within the fourth immunoglobulin-like domain, while the other variant in the Golden Retriever mix is further downstream within the protein kinase domain (**Fig 4C**).

Both mutations are splice-acceptors, making it difficult to determine their impact on transcription. The affected exons may be skipped entirely, or an alternative splice site may be used, resulting in either partial exon retention or retention of nearby intronic sequence. Dogs carrying the *PDGFRA* mutations had no reported health concerns, which is surprising as *PDGFRA* is a broadly expressed developmental gene and strongly associated with gastrointestinal stromal tumor-plus syndrome in humans (Manley et al. 2018; Huang et al. 2022).

Two other breeds share a similar phenotype as the Çatalburun, the Andean tiger hound, about which little is known, and the Spanish Pachón Navarro. The Pachón Navarro, like the Çatalburun, is a pointing breed, and not all breed members have a bifid nose. Evidence for bifid nose dogs in Spain can be observed not only in 18^th^ century architecture (**Fig. 4D**), but also in a 1623 painting of a hunting party titled “Diane et ses nymphes s’apprêtant à partir pour la chasse” by Pierre Paul Rubens and Jan Brueghel l’Ancien. Other works from around the turn of the 20^th^ century describe bifid nose pointing dogs throughout the south of France (Arkwright 1906). Although it is unknown whether the bifid nose phenotype observed in the Çatalburun and Pachón Navarro breeds has a common origin, Turkey and Spain were both under the rule of the Umayyad Caliphate in the 8^th^ century CE, providing a possible route for cultural exchange between the two regions. Given our observation of multiple variant alleles among bifid dogs recruited for this study and the recurrent artistic depictions, the phenotype could have arisen multiple times in pointers.

The newly discovered *LCORL* c.1978_1981delACAG allele was identified because of its high frequency in Spanish Mastiffs, Berger Picards, and Briards (**Table 1**). Like the earlier discovered *LCORL* c.3768_2769insA variant, the c.1978_1981delACAG allele causes a frameshift and premature stop, resulting in loss of the DUF4553 DNA-binding domain from the long isoform of LCORL (Plassais et al. 2019). Both alleles are associated with larger size and stature in dogs (**Fig 4F,G**). *LCORL* is also a primary determinant of stature across many domesticated mammals including goats (Saif et al. 2020; Graber et al. 2022), cattle (Bouwman et al. 2018; Chen et al. 2020), pigs (Rubin et al. 2012), and horses (Metzger et al. 2013; Tetens et al. 2013). Given that the gene is under selection in multiple species, it is unsurprising that more than one *LCORL* allele is under selection. The presence of both alleles within the same breeds also suggests that both variants predate breed development. Further investigation into the mechanisms by which trait variants affect phenotypes may reveal other instances where selection for different alleles achieves redundancy, thus assuring the same phenotypic outcome.

The association between breed-differentiated regions and potential trait variants in constrained genes returned few associated CDS variants. Importantly, none of these associations included OMIA variants prioritized using genotype-first approaches, indicating that signatures for breed enrichment are distinct from signatures for selection/breed differentiation.

This is because many OMIA variants are disease alleles and, while overrepresented in a small subset of breeds, are not under positive selection (Boyko et al. 2010; Kim et al. 2018; Plassais et al. 2019). With additional disease risk phenotyping for dog breeds, genotype-first approaches may be useful for expanding lists for disease risk alleles. Most genes in breed-differentiated regions had no association with potentially functional CDS variants, consistent with non-coding variation being the primary driver of phenotypic variation across breeds, a repeated finding from other large-scale multi-breed canine genetic analyses in dogs (Sahlen et al. 2021; Dutrow et al. 2022; Morrill et al. 2022) and humans (Halldorsson et al. 2022). However, breed-differentiated regions were enriched for constrained genes, emphasizing the importance of constraint in predicting phenotypic impacts.

One of the defining features of the canine system is that often few loci contribute to large phenotypic variation, particularly for morphological traits like breed standard weight and height or ear shape and position (Rimbault et al. 2013; Plassais et al. 2019), while in the human genome there are often large numbers of loci associated with the same phenotypes (Yengo et al. 2022). Loci contributing to extreme phenotypic differences between breeds overwhelmingly exhibit strong signatures of positive selection because of artificial selection for desired traits (Plassais et al. 2019; Morrill et al. 2022; Plassais et al. 2022). Our analysis shows that highly constrained genes are overrepresented in such regions and are therefore the likely targets of trait selection, driving phenotypic variation. This suggests an explanation for why CDS variation is rarely associated with morphological differences between breeds and more often associated with disease. Constrained genes are often essential for development and survival and are frequently implicated in highly penetrant genetic diseases and syndromes (Lek et al. 2016; Karczewski et al. 2020; Sun et al. 2024; Zeng et al. 2024). Trait selection in dogs is therefore a balancing act between preserving essential gene functions and maximizing regulatory variation of essential genes to drive phenotypic extremes. Since breed variation is driven by the expression of genes with essential biological roles, future work should focus on assigning non- coding variants under positive selection to their target genes.

Central to the analysis is the identification of variants with phenotypic impacts, which was assessed according to evolutionary/functional constraint acting on genes/sites containing the variant in question. However, specific factors impact the interpretation of these results. For instance, the measured selection effects only capture negative selection acting on heterozygous variants with highly deleterious impacts on fitness (Fuller et al. 2019). Our analysis is therefore blind to recessive deleterious alleles and genes associated with late age of onset conditions such as breed-prevalent diseases like cancer. Since singletons are the input to the mutation model, depletion of variants with little to no deleterious impact in a heterozygous state remain unobservable, as do variants that produce no phenotype until after a dog’s reproductive years have elapsed. This is relevant to canines where a high degree of inbreeding makes dogs especially susceptible to genetic load and recessive disease allele burden (Marsden et al. 2016; Mooney et al. 2021; Donner et al. 2023). Moreover, OMIA variants segregating in canine populations likely persist because they do not have strong effects on fitness in a heterozygous state. In fact, many of the OMIA variants underlie traits, such as coat color or recessive and late- onset diseases, such as the many forms of progressive retinal atrophy (Miyadera et al. 2012; Downs et al. 2014; Bunel et al. 2019). Therefore, strong human ortholog LOEUF constraint lacks an association with OMIA variants because OMIA LoF variants have a low impact on fitness, and human ortholog LOEUF constraint indicates genes where heterozygous LoF mutations have a strong deleterious impact on fitness. Another factor they may impact interpretation of results is whether the unit for measuring constraint is gene or base-pair. The increased resolution provided by site-specific phyloP scores is only beneficial for analyzing missense mutations as mutations annotated as LoF, defined throughout as premature stop codons, frameshifts, and splice-site disruptions, often impact the entire transcript downstream of the mutation. Missense mutations, alternatively, have more localized effects and usually cause negligible impacts on biology unless interrupting important functional domains (Gerasimavicius et al. 2022). Therefore, in dogs, human ortholog LOEUF is the best of the three constraint metrics for prioritizing LoF mutations for phenotypes with severe impacts on fitness. Conversely, variant phyloP scores are best for prioritizing missense mutations.

While the genotype first approach is ideal for a species with a complicated population structure like dogs, it is facilitated by sufficient breed representation and availability of accurate phenotype information. While mapping breed standard traits has proven fruitful, more refined phenotyping and mapping of more nuanced traits will require individualized measures of each dog. For instance, CT scans are now fundamental to genetic studies of canine skull morphology, as reliance on owner measurement and breed descriptors lacks precision (Marchant et al. 2017).

The dog genome project began with two goals, and both remain a high priority today: improve the health of canine companions through informed genetics and translate those findings to advance human health. While many breeds feature largely healthy, long-lived animals, others are characterized by an excess of disease alleles and associated health problems (Axelsson et al. 2021). These breeds offer a unique opportunity to make use of the approaches and data provided here to identify underlying genetic flaws, particularly for diseases of relevance to human health. Of equal priority are studies of morphologic and behavioral genetics, as identification of alleles that control extreme phenotypes associated with these traits can improve our understanding of human growth and development.

## Methods

### Defining a canine mutation model and callable gene set

We built a mutation model in dogs following the methods described in (Samocha et al. 2014). Briefly, trinucleotide substitution rates, normalized according to the generation mutation rate and trinucleotide frequencies, were applied to each possible substitution for each site within gene models. For each gene model, substitution probabilities were summed according to their consequences on coding sequence, such as synonymous, missense, stop gain, stop loss, etcetera. The result was per gene probabilities for each mutation type per generation (**Supplemental Fig. S4**). Dog10K variants at least 20 bp from an exon and within callable loci were used to calculate trinucleotide substation rates (Meadows et al. 2023). A canine generation mutation rate of 9.02x10^-9^ was used for normalization (Bergeron et al. 2023) (**Supplemental Fig. S5**). We calculated transcript-specific mutation probabilities using NCBI gene annotations from Dog10K, only considering sites that were callable loci, obviating the need to normalize probabilities by mean exon coverage (**Supplemental Table S15**). There was insufficient variant data to adjust mutation probabilities by evolutionary conservation as was performed in (Samocha et al. 2014). We classified genes according to intact start and stop codons, low-quality annotation, and partial annotation. Also, for each gene, a representative transcript was chosen based on reciprocal best hit blast results with protein sequences of human representative transcripts. Where no blast hit was found, the first annotated transcript was used as a representative. Blast hits were also defined as dog-human orthologs and used to assign human LOEUF constraint metrics to dog genes. For phyloP constraint, we used the Zoonomia values mapped to UU_Cfam_GSD_1.0 as part of Dog10K, which were further summarized according to gene models (**Supplemental Table S16**). Expected mutations per gene were calculated by sampling *n* gene names with replacement weighted by gene/gene group mutation probabilities, where *n* is the number of observations the expected rate is being compared to. Sampled gene names are tabulated to get the expected per gene mutation count. This process is replicated 10,000 times to generate 95% expected ranges.

### Preparing genotypes for analysis

Genotyping and filtering procedures are described in full in the supplemental methods. Briefly, a VCF comprised of SRA genomes was constructed using GATK to complement the publicly available Dog10K VCF (Poplin et al. 2017). All WGS data was mapped to the UU_Cfam_GSD_1.0 reference assembly. For each VCF, samples were filtered according to a variety of genotype metrics (**Supplemental Table S17**), which included coverage statistics (**Supplemental Fig. S6**), heterozygous allele balance (**Supplemental Fig. S7**), as well as several others (**Supplemental Fig. S8**). A total of 1,146 SRA and 1,778 Dog10K samples remained (**Supplemental Table S18**). Duplicates and first-degree relatives across both datasets were removed from the analysis (**Supplemental Fig. S9**). A total of 2,291 samples remained (**Supplemental Table S19**). Sites were filtered according to variant quality score recalibration (VQRS) using a common training set (**Supplemental Fig. S10**). Autosome and chromosome X PAR sites (**Supplemental Table S20**) were analyzed separately from chromosome X non-PAR sites (**Supplemental Table S21**). Sites that passed VQSR filtering in both datasets were used to create a merged VCF for population-scale analyses. Sites that passed filtering in either dataset and were within the CDS underwent further filtering (**Supplemental Fig. S11**) (**Supplemental Table S22**). Criteria for filtering CDS genotypes depended on chromosome, sex, and genotype (**Supplemental Table S23**). CDS indels and SNVs were reclassified and atomized in both datasets to improve harmonization (**Supplemental Fig. S12**) (**Supplemental Table S24**). Over 95% of SNVs and over 80% of indels passed filtering (**Supplemental Table S25**). Genotype quality was the criteria that failed most often (**Supplemental Table S26**). Sample genotype pass rates were higher for SNVs than for indels (**Supplemental Table S27**). CDS filtering removed genotyping biases between each dataset for SNVs (**Supplemental Fig. S13**) and indels (**Supplemental Fig. S14**). Genotypes were merged and annotated with variant effect predictor (VEP) (McLaren et al. 2016). All VCF-based operations were performed with bcftools (Danecek et al. 2021), plink (Purcell et al. 2007), vcftools (Danecek et al. 2011), and vcflib (Garrison et al. 2022).

### Defining breeds and groups

Illumina CanineHD array genotypes from 2,156 samples were lifted over to the UU_Cfam_GSD_1.0 reference genome using LiftoverVCF (**Supplemental Table S28**). WGS sites were downsampled to match array positions and datasets were merged. After filtering at MAF > 0.01, genotyping rate > 0.9, and LD < 0.5, a total of 76,570 sites remained (Purcell et al. 2007). Using Phylip, a consensus neighbor joining tree was built with Golden Jackal set as the outgroup (Felsenstein 1993). Branch support was calculated from 10% jackknifing 10,000 times. Branches needed at least 50% support. Breed/group clusters were determined by optimizing for shared breed membership on the same branch. Samples that did not match the majority breed on a branch were assigned new breed membership, matching the majority breed. Branches that consisted of overlapping breeds were merged. Breeds that could not form a branch of at least three members were placed in the unassigned category.

### Canine population statistics and breed differentiated loci

Population analyses were only performed on the 200 breeds/groups that had five or more members. Genome-wide population statistics were calculated across 200,000 randomly sampled sites. Nucleotide diversity within breeds was measured as pi, the average distance between two randomly sampled sequences in a breed (Nei and Li 1979). Breed differentiation was measured as F_ST_, using the Weir and Cockerham estimator on a per-site basis (Weir and Cockerham 1984). F_ROH_ was calculated using the same procedures as specified in (Meadows et al. 2023). We detected breed-differentiated regions by calculating F_ST_ between all 200 pairs of breeds with five or more members across 50kb windows using a 10kb step size. Windows with F_ST_ values > 0.8, which corresponded to ∼0.1% of all pairwise comparisons, were extracted for further analysis. The density of high F_ST_ windows was then measured across the genome and the “peaks” function of IDPmisc, a peak-calling software R package, was used to define breed differentiated regions (Locher 2024). Next, for each region, we extracted breeds with pairwise F_ST_ values > 0.8 within the region’s peak window. Extracted breed pairs with F_ST_ < 0.05 were defined as sharing the same haplotype and were assigned the same cluster ID. We also defined genomic segments of extended breed homozygosity if half of the 50kb windows along a megabase segment had pi values < 0.02. Breed-differentiated regions were defined as selection loci in individuals where there was overlap with extended homozygosity regions.

### Identification of additional orofacial cleft genes

Blood or saliva samples were collected from nine additional dogs, eight with a bifid nose, of which two were littermates. The non-bifid nose dog is a parent to the littermates. Samples were whole genome sequenced at mean 20x coverage and genotyped following the above methods used for SRA samples. Variant sites shared with the Dog10K cohort were filtered out from the analysis. Next, SVNs were discarded if QD < 2, SOR > 3, FS > 60, MQ < 40, MQRankSum < - 12.5, or ReadPosRankSum < -8.0. Indels were discarded if QD < 2, FS > 200, ReadPosRankSum < -20.0, or variant length was > 10 bp. Since orofacial clefts within this cohort were likely sporadic, except for the siblings and parent, we focused specifically on high quality heterozygous genotypes that met the following criteria: GT = 0/1, GQ >= 20, DP >= 10, AD ratio > 0.5, and AD ratio < 2.0. Next, we selected variants as candidates if they were found in any orofacial cleft candidate genes, which were determined according to membership within the following gene lists; DisGeNET associations for craniofacial abnormalities (C0376634), cleft Palate (C0008925), or cleft upper lip (C0008924) (Pinero et al. 2017), experimentally identified orofacial cleft genes (Xu et al. 2021), and genes with restricted craniofacial expression during human development (Yankee et al. 2023).

## Acknowledgements

We thank Tatiana Feuerborn and Heidi Parker for their helpful advice and insight throughout the project and Gabby Spatola for assistance in recruiting samples. We also thank Zachary Lounsberry and Embark Veterinary for assistance in recruiting samples and thank Sruthi Hundi for bioinformatic assistance. For careful reading of the manuscript and providing helpful feedback, we also thank Christopher B. Kaelin and Wesley C. Warren. Most importantly, we thank all citizen scientists who donated samples for analysis, without whom this work would be impossible. E.A.O., R.M.B., A.C.H., D.L.D., A.S.A are funded by the Intramural Program of the National Human Genome Research Institute at the National Institutes of Health (HG200377). H.L. is funded by the Jane and Aatos Erkko Foundation.

## Competing interest statement

A.R.B. is a co-founder and board member of Embark Veterinary, Inc., a private canine DNA testing company. All other authors declare no competing interests.

**CRediT author statement** Conceptualization: R.M.B. and E.A.O. Methodology: R.M.B. and A.S.A. Software: R.M.B. and A.S.A. Validation: N.B., F.G.V.S., and M.K.H. Formal analysis: R.M.B.

Investigation: R.M.B., N.B., M.K.H., and D.L.D.

Resources: N.B., P.S., C.T., M.E., F.G.V.S., M.K.H., J.H., H.L., B.C.K., A.R.B., D.L.D., and R.M.B.

Data Curation: R.M.B. and A.C.H.

Writing - Original Draft: R.M.B. and E.A.O.

Writing - Review & Editing: R.M.B., E.A.O., and D.L.D. Visualization: R.M.B.

Supervision: E.A.O.

Project administration: R.M.B. and E.A.O.

Funding acquisition: E.A.O.

