## Supplemental Methods for "A genotype-first approach identifies variants for orofacial clefts and other phenotypes in dogs"

**Supplementary methods**

**Constructing a high-quality cohort for analysis of canine coding variation**

To investigate genetic variation within the coding regions of dogs, we assembled one of the largest high-quality WGS canine datasets to date. We merged the VCF from the Dog10K project, a multi-institutional effort to sequence thousands of dogs using a unified approach, and a VCF made from dog genomes from the Short Read Archive (SRA), named throughout as the Dog10K VCF and SRA VCF, respectively (Meadows et al. 2023). There are several key challenges inherent in the process of merging and harmonizing data from a variety of sources. These include selection of high-quality samples, differences in genotyping software, selection of representative sites, and identification of closely related/duplicated samples.

*Read alignment, data processing, and genotyping*

Read alignment and genotyping were performed independently for Dog10K and SRA. Documentation for Dog10K data processing is provided in Meadows et al. (2022). Both Dog10K and SRA used the same reference genome, which was a combination of autosomes and chromosome X from UU_Cfam_GSD_1.0 and chromosome Y sequence from ROS_Cfam_1.0 (Meadows et al. 2023). Genotypes were called for chromosomes 1 through X, while Y sequences were included to prevent mapping of Y chromosome reads to other chromosomes. Importantly, each dataset was processed using different versions of the Genome Analysis Toolkit (GATK). Dog10K was processed with GATK version 4.2.0.0 and SRA was processed with GATK version 4.4.0.0. One of the key differences between these software versions is that as of version 4.2.3.0 the symbol for missing genotypes, “./.”, was altered to the homozygous reference genotype, “0/0”, where missing calls are identified by a genotyping depth (DP) of 0. Another difference is that each treated chromosome X calls in male dogs differently. In Dog10K males were genotyped as haploid and in the SRA VCF males were genotyped as diploid. These specific differences between the two datasets were carefully considered when performing sample filtering, site filtering, and VCF merging on both autosomes and sex chromosomes.

*Sample filtering*

Samples were selected based on VCF genotype level data from each of the Dog10K and SRA datasets. Specific genotype statistics collected include genotype (GT), depth (DP), allele depth (AD), and genotype quality (GQ). These values were only collected from sites that passed initial VQSR filtering (99^th^ percentile truth sensitivity) and from genotype fields where the GT value was either “0/0”, “0/1”, “1/1”, or “./.”. Samples with mean DP below 15 were removed first (**Supplemental Fig. S7**). Next, we counted the total number of genotyped sites per individual. Samples were kept if the total number of genotypes was within five standard deviations of the mean number of genotypes. Any samples that had no genotypes within a 5Mb segment were also removed. We also defined pass and fail criteria for each genotype statistic according to GT state (**Supplemental Table S18**). Samples with greater than a 5% fail rate for any of the defined criteria were removed from the analysis. To determine an appropriate allele balance cutoff, we measured the proportion of samples that would have heterozygous fail rates > 5% at different cutoff values (**Supplemental Fig. S8**). We found that by using an allele balance cutoff of 20%, we were able to remove only a small fraction of samples. Allele balance cutoff values of 25% led to most samples within Dog10K and ~40% of SRA samples having failure rates > 5%. Altogether, 1,165 SRA samples and 1,778 Dog10K samples passed all initial filters (**Supplemental Table 19**). Samples with coverage < 20X were particularly susceptible to failure, however, and many samples at ~ 20x mean coverage failed to have consistent coverage of > 10x across the whole genome (**Supplemental Fig. S9**). Importantly, 165 SRA samples and 195 Dog10K samples were eliminated for failing only one of the filtering criteria, indicating the importance of using a diverse array of sample filters.

To identify a set of unique unrelated samples across both datasets, we analyzed kinship coefficients between each pair of samples. Specifically, we extracted canine axiom array sites from the SRA VCF and Dog10K VCF. We then merged both VCFs containing the extracted genotypes and removed sites with allele frequencies less than 5%. We then used plink2 to calculate pairwise kinship coefficients using the “--make-king-table” function (Manichaikul et al. 2010). Once relatedness values were determined we identified duplicated samples as sharing a kinship coefficient > 0.45. For every pair of duplicated samples, we removed one sample until no more duplicates were present in the dataset. If duplicate samples were found across both datasets, we removed the SRA sample and kept the Dog10K sample. Next, we identified first-degree relatives as sharing a kinship coefficient between $\frac{1}{2^{5/2}}$ and $\frac{1}{2^{3/2}}$ and we flagged sample relationships with proportion of IBS0 < 0.005 (**Supplemental Fig. S10**). The number of first-degree relatives each sample shared was calculated and the sample sharing the most first-degree relatives was removed. This process of removing the most highly related sample was repeated until no first-degree relatives remained in the dataset. After duplicate and first-degree relative removal, 572 samples remained in the SRA dataset and 1,719 samples remained in the Dog10K dataset (**Supplemental Table S20**). After filtering, the merged dataset between SRA and Dog10K consists of 2,291 unique, high-quality samples with no siblings or parent-offspring pairs.

*Initial site filtering*

An important goal of this analysis is to increase representation of canine genetic diversity by merging genetic data from two different VCF files. Critical to this process is harmonization of filtering procedures across both datasets, as removal of sites that were flagged as low quality in one dataset and as high quality in the other dataset can create batch effects and lead to incorrect allele frequency estimates. Here we describe our approach for harmonizing site-based filtering in the SRA and Dog10K datasets.

To harmonize and filter SNV sites from the SRA dataset and Dog10K dataset, we performed VQSR for each VCF using a common training/truth marker set, overwriting previous VQSR annotations. For autosomal + chrX PAR data, we selected a truth sensitivity threshold of 99.7 and for chrX non-PAR data, we selected a truth sensitivity threshold of 99.5. We selected these threshold values as they each contained the greatest number of sites for both SRA and Dog10K while also maintaining a higher tranche specific true positives rate than tranche specific false positive rate (**Supplemental Fig. S11**). As expected, the SRA VCF contained a higher fraction of false positive sites than the Dog10K VCF. This is likely because the SRA VCF contained many low-coverage samples that each contributed low-confidence sample specific variants. For autosomes + chrX PAR, the SRA dataset contained ~47M PASS sites with a Ti/Tv ratio of 2.016 and the Dog10K dataset contained ~44M PASS sites with a Ti/Tv ratio of 2.1005 (**Supplemental** **Table S21**). For chrX non-PAR, the SRA dataset contained ~1.4M PASS sites with a Ti/Tv ratio of 1.675 and the Dog10K dataset contained ~1.3M PASS sites with a Ti/Tv ratio of 1.7425 (**Supplemental** **Table S22**). To perform filtering on indels, we used hard thresholding of site-specific metrics to remove low quality sites. These threshold values were “QD < 2.0”, “FS > 200.0”, “ReadPosRankSum < -2.0”, and “SOR > 10.0”.

Next, we removed all samples that failed filtering and relatedness criteria described above from the SRA and Dog10K datasets. We then removed any site from each dataset that had an updated allele count (AC) of 0. The bcftools isec function was then used to compare filter status of all remaining sites in SRA and Dog10K across the whole genome and the coding sequences (**Supplemental Fig. S12**). Importantly, we found that between 1.99% and 4.92% of sites that passed filtering in one dataset had failed flittering in the other dataset, indicating the importance of comparing filter statuses of sites prior to merging VCF files (**Supplemental Table S23**). Based on this result, we used two different approaches to select sites that would facilitate VCF merging in downstream analyses. The first approach is for whole genome population-based analyses, where rare variants are often removed and SNVs are treated as markers to measure genetic differences and similarities between individuals and groups. Here, we only select SNVs that PASS filtering criteria in both VCF files. This approach helps ensure that likely common variants that are consistently identified as high-quality are used to calculate population-based statistics. Altogether we select ~ 22.2M SNVs for whole genome population genetic analyses. The second approach is for capturing variants with potential functional impacts on gene sequence. In this case we select SNVs and indels within the coding sequence that pass filtering in either dataset. It is important to capture rare variants that may only be found in one dog and not shared between both datasets, as variants with large phenotypic impacts tend to be rare. All together we selected ~ 821k SNV sites and ~ 70k indel sites from within the coding regions of the dog genome for further analysis.

*Filtering CDS sites*

To determine appropriate filtering thresholds for genotypes we analyzed the distributions of genotype statistics for different sets of SNV and indel sites selected at random. These sets included 5000 SNV sites from Autosomes + chrX PAR, 1000 SNV sites from chrX non-PAR, 1000 indel sites from Autosomes + chrX PAR, and 500 indel sites from chrX non-PAR. Multiallelic sites were then split into separate rows and the following genotype values were extracted: GT, AD, DP, and GQ, corresponding to genotype, allele depth, depth, and genotype quality, respectively. This task was performed using the vcflib v1.0.3 vcf2tsv function with the “-g” flag present, which converts VCF files and genotypes into long format (Garrison et al. 2022). Genotypes were analyzed according to whether they were homozygous reference (ref), heterozygous (het), or homozygous alternate (alt). In the case of males on the non-PAR region of chromosome X, genotypes were considered as either reference (ref) or alternate (alt), due to their haploid nature. Allele balance was calculated as the alternate read count from AD (A2) divided by the value for DP. The genotype filtering thresholds that were ultimately used are displayed in **Supplemental Table S24**.

We found in many cases SNVs and indels map to the same position, creating multi-allelic variants. This caused a problem in our analysis, as the SNV at the multiallelic site ended up being classified as an indel. To overcome this, we reclassified SNVs and Indels. For both variant types we split multiallelic variants with “bcftoools norm -m –“ and atomized them with “bcftools norm -a”. Next. we pooled all variants and then once again separated SNVs and indels. We tested the outcome of this procedure by measuring the number of high frequency variants in one dataset that have an allele count of 0 in the other dataset (**Supplemental Fig S13**). After reclassifying indels, the number of dataset-specific high frequency variants decreased by an order of magnitude (**Supplemental Table S25**).

Next, we applied filtering procedures to the genotype data for each dataset. Genotypes that failed any of the criteria were set to missing. All together, we retrained over 95% of SNV genotypes and for indels, we retained over 81% autosomal and over 88% chromosome X genotypes (**Supplemental Table S26**). Minimum GQ was the filtering criteria that was most frequently failed with > 4% of indel genotypes failing allele balance thresholds (**Supplemental Table S27**). The majority of samples had SNV genotyping pass rates > 95% and indel genotyping pass rates > 80% (**Supplemental Table S28**).

Finally, we tested the outcomes of genotype filtering for removing dataset-specific bias. For Dog10K and SRA, we compared the distributions of variants per individual. For SNVs and indels, in both males and females, across the autosome and chromosome X, genotype filtering caused a large decrease in the distance between each distribution, indicating that filtering procedures remove a high degree of dataset-specific bias (**supplemental Fig. S14**) (**Supplemental Fig. S15**).

**Primer sequences used for variant validation**

Çatalburun - chr13:47440160, A/C, PDGFRA, splice acceptor

Fwd: 5` - TGTTCTGGCCACATAAGCTG – 3’

Rev: 5` - TTCACTTCATCACCGAGCTG – 3’

Newfoundland dog DCM - chr28:5754409, CGTTAC/CTC , ANKRD1, inframe indel

Fwd: 5’- CTCCTCCAACGCTGAAATCCT -3’

Rev: 5’- CAAAGACCGAGTGAGTACCT -3’
