## Supplemental Figures for "A genotype-first approach identifies variants for orofacial clefts and other phenotypes in dogs"

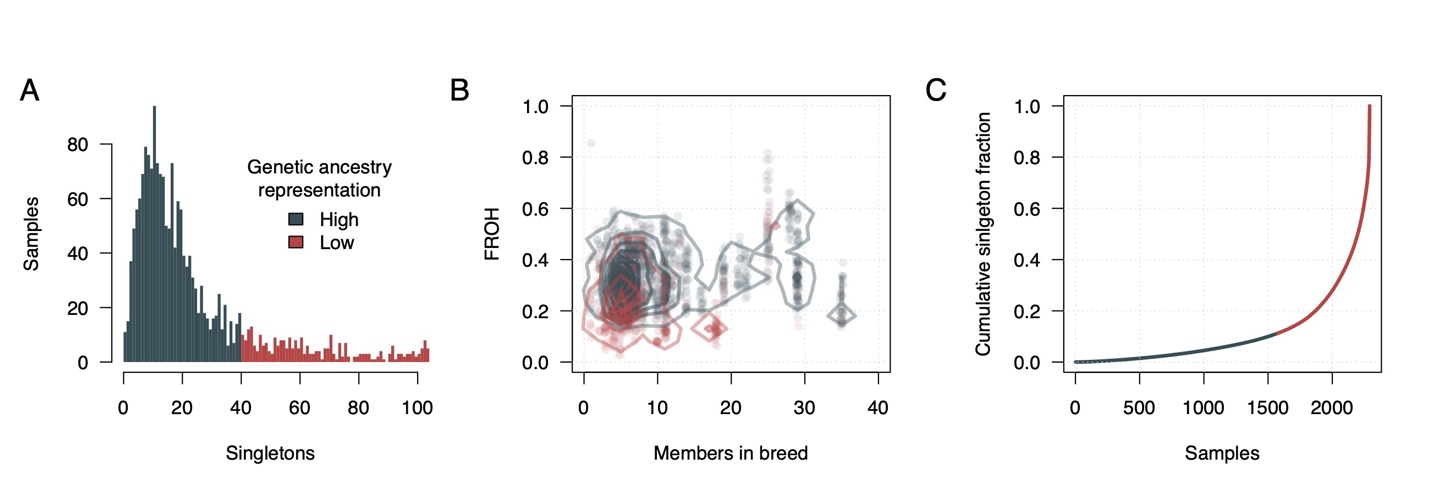

**Supplemental Fig. S1: Characterizing genetic ancestry representation in dogs.** A) Singletons per dog. Dogs with less than 40 singletons were classified as having highly represented ancestry. B) Dogs with highly represented ancestry have more breed members in the dataset and higher rates of F_ROH_ than dogs with low ancestry represented ancestry. C) A minority of singletons are found in dogs with highly represented ancestry.

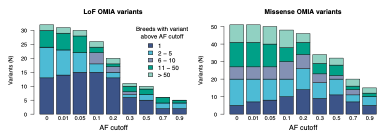

**Supplemental Fig. S2: Breed specificity of OMIA variants within the analysis cohort**

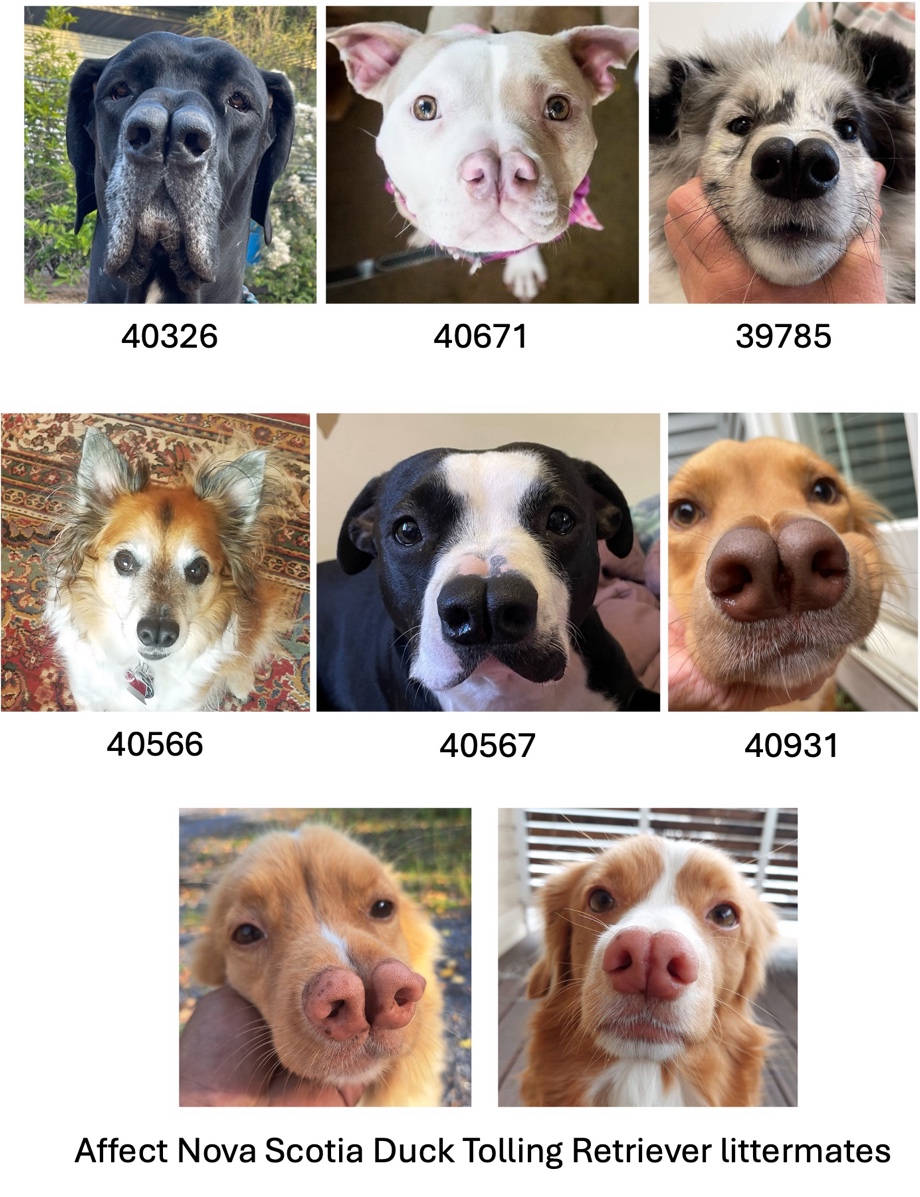

**Supplemental Fig. S3: Recruited dogs with bifid nose phenotype**

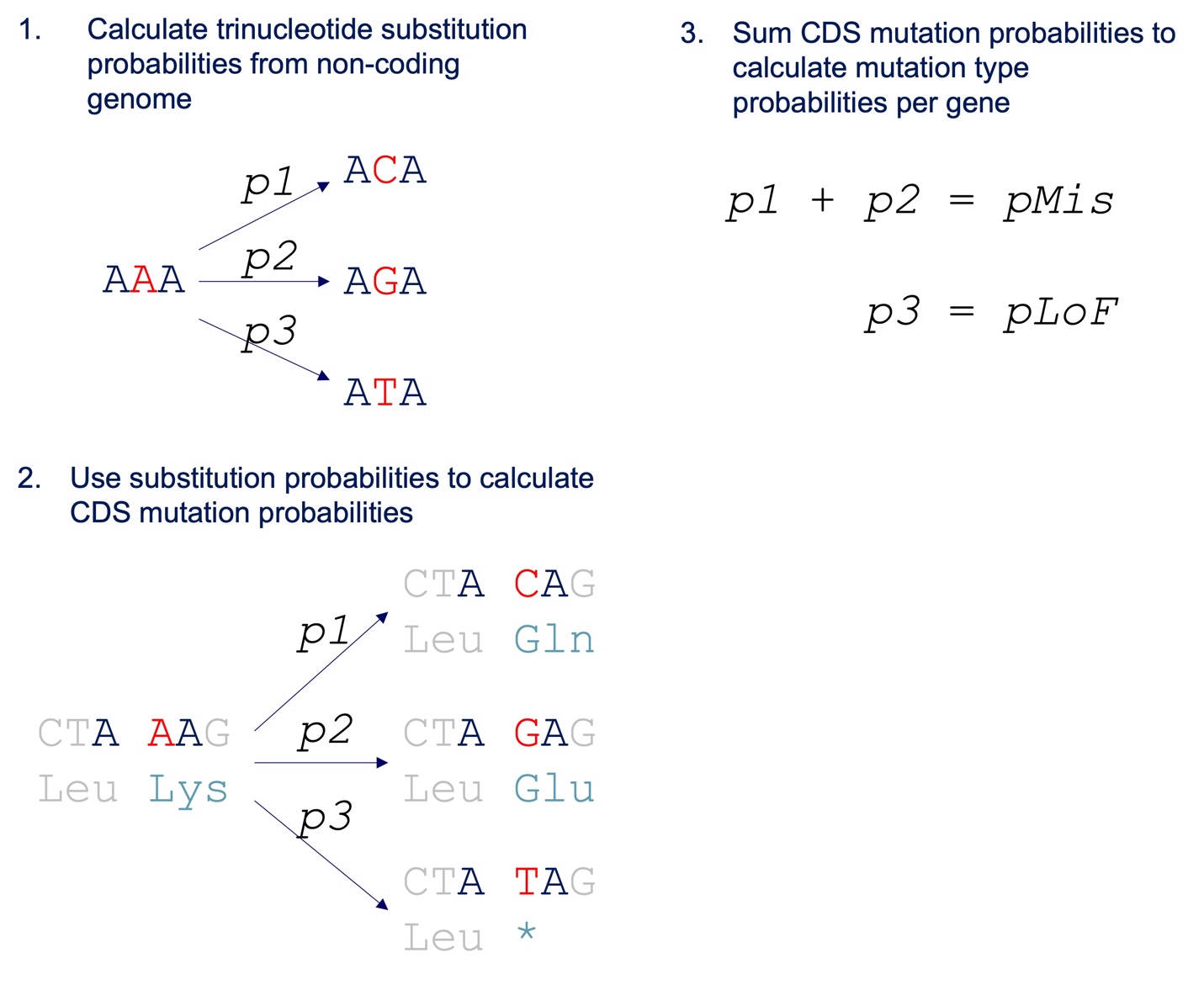

**Supplemental Fig S4: Steps for building mutation model.**

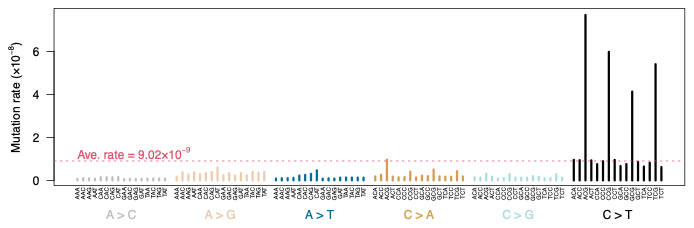

**Supplemental Fig S5: Trinucleotide mutation rates in dogs.**

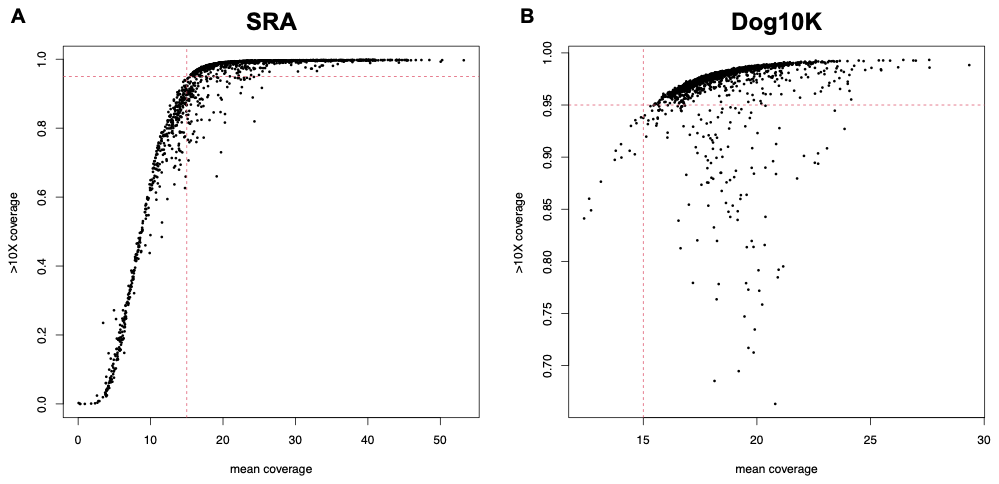

**Supplemental Fig. S6: mean coverage of sites and proportion of sites with coverage > 10x for all samples.** A) Results for SRA dataset. B) Results for Dog10K dataset. Red dotted lines in A and B indicate minimum threshold values for samples to pass filtering criteria.

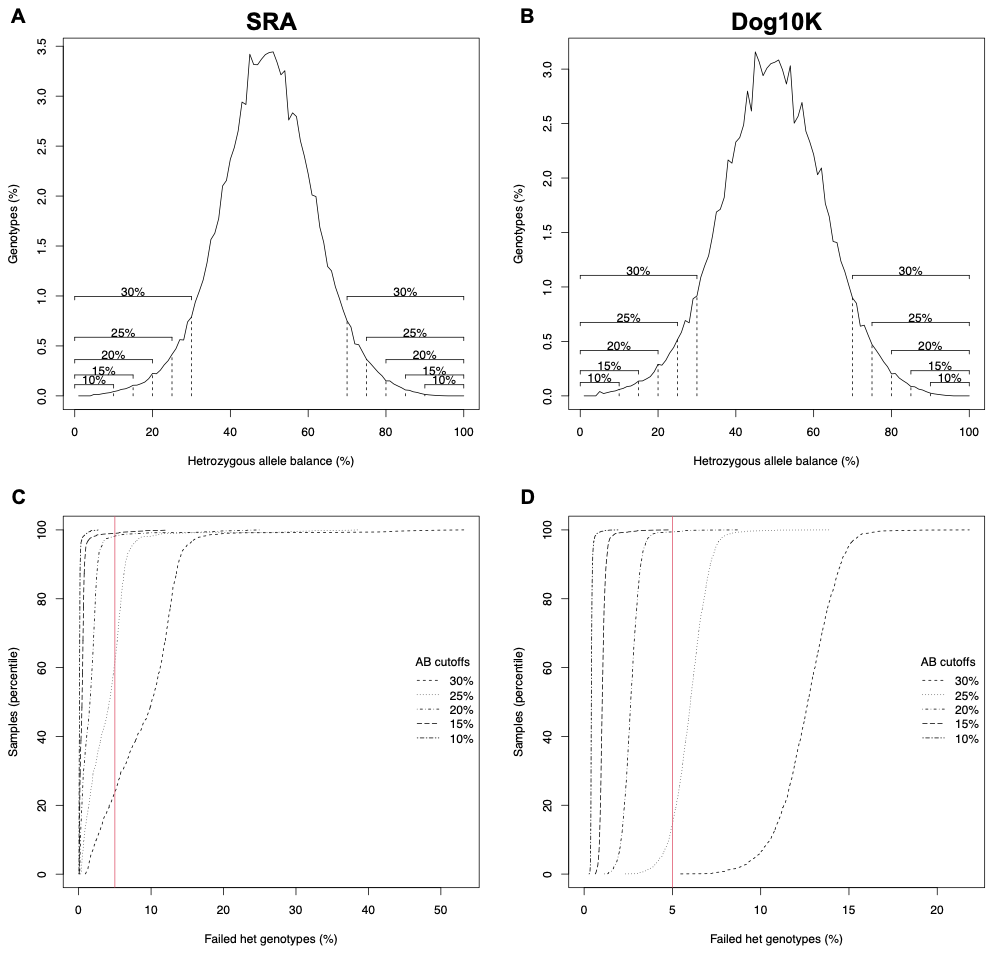

**Supplemental Fig. S7: Distributions of allele balance for heterozygous genotypes.** A-B) Allele balance (AB) for all heterozygous genotypes in the A) SRA dataset and the B) Dog10K dataset. Vertical dotted lines indicate different allele balance cutoff values for declaring failed het genotypes. C-D) The cumulative distribution of per sample genotype failure rates at each allele balance cutoff for the C) SRA dataset and the D) Dog10K dataset. The red solid line indicates a per sample heterozygous failure rate of 5%. For example, in D, All samples have > 5% heterozygous failure rate at an AB cutoff of 30%.

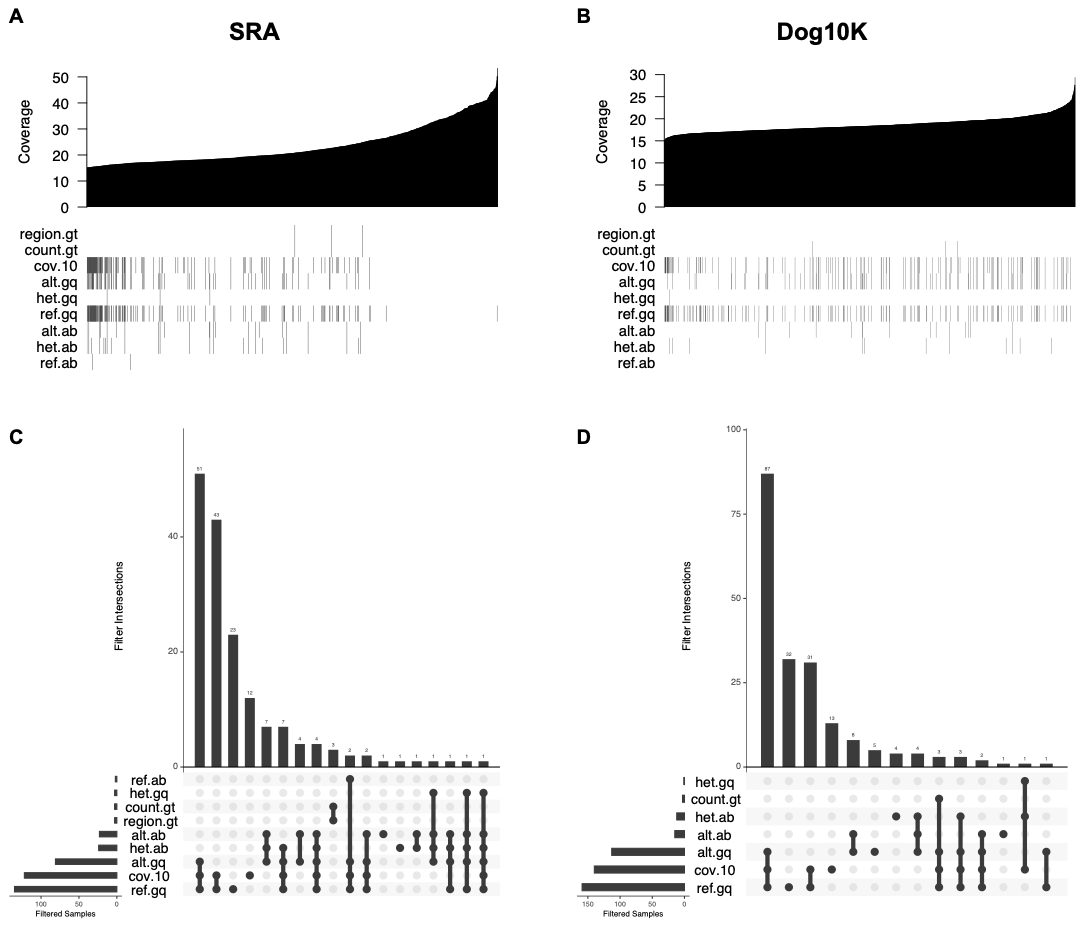

**Supplemental Fig. S8: Samples removed for failing filtering criteria at rates > 5%.** Filtering criteria are described in table X. A-B) Samples are sorted along the X-axis according to mean coverage level across all sites. Shading is used to indicate samples in the A) SRA dataset or the B) Dog10K dataset that were removed for failing a specific criterion. C-D) Upset plots indicating the number of samples removed for failing different combinations of filtering criteria. Data is provided for both the C) SRA dataset and the D) Dog10K dataset.

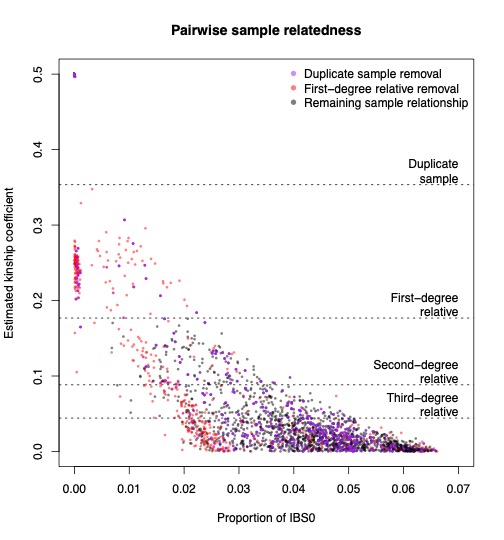

**Supplemental Fig. S9: Relatedness of remaining samples after quality filtering and merging.** The estimated kinship coefficient reflects the degree of relatedness between two samples. The proportion of IBS0 indicates the fraction of tested sites where no identity by state was observed between a pair of samples. Each datapoint indicates a pairwise relationship between two samples. Purple datapoints were removed after filtering out duplicated samples. Red datapoints were removed after iteratively filtering out the samples with the most first-degree relatives. The black datapoints remain after all filtering and have kinship coefficient values below the first-degree relative cutoff.

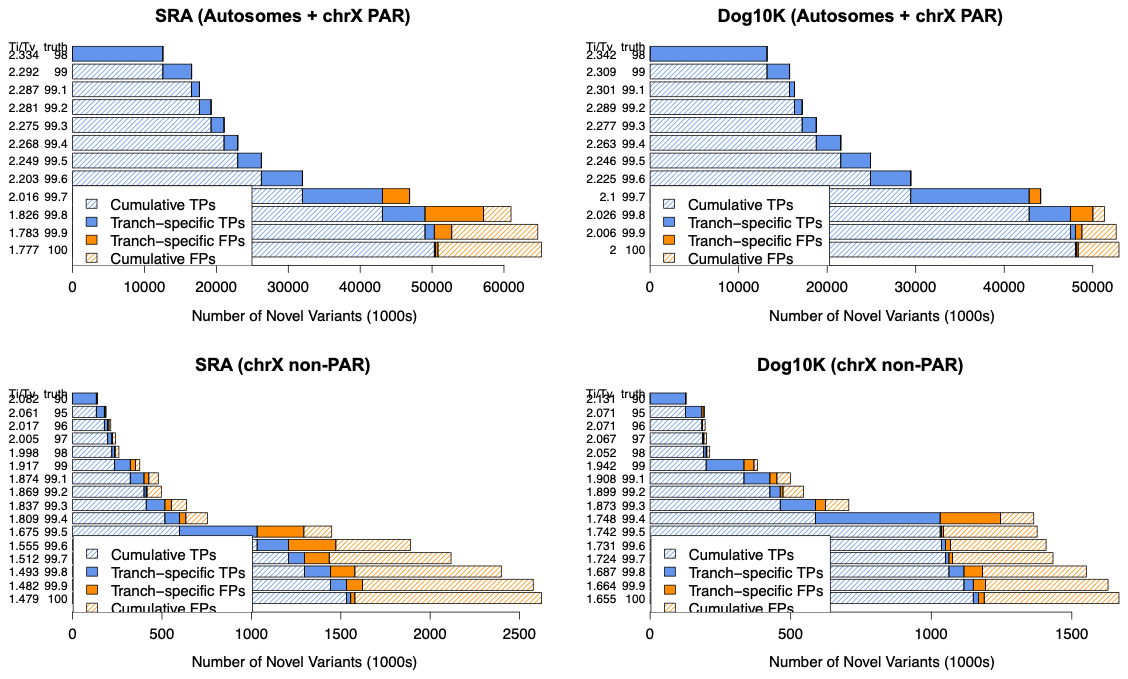

**Supplemental Fig. S10: Graphical output from VariantRecalibrator (GATK v4.5.0.0) for the SRA and Dog10K datasets and both autosomes and the X chromosome.** At each truth sensitivity threshold, the Ti/Tv ratio, the total number of variants within each tranche, and the proportion of true positive (TP) and false positive (FP) variants are displayed.

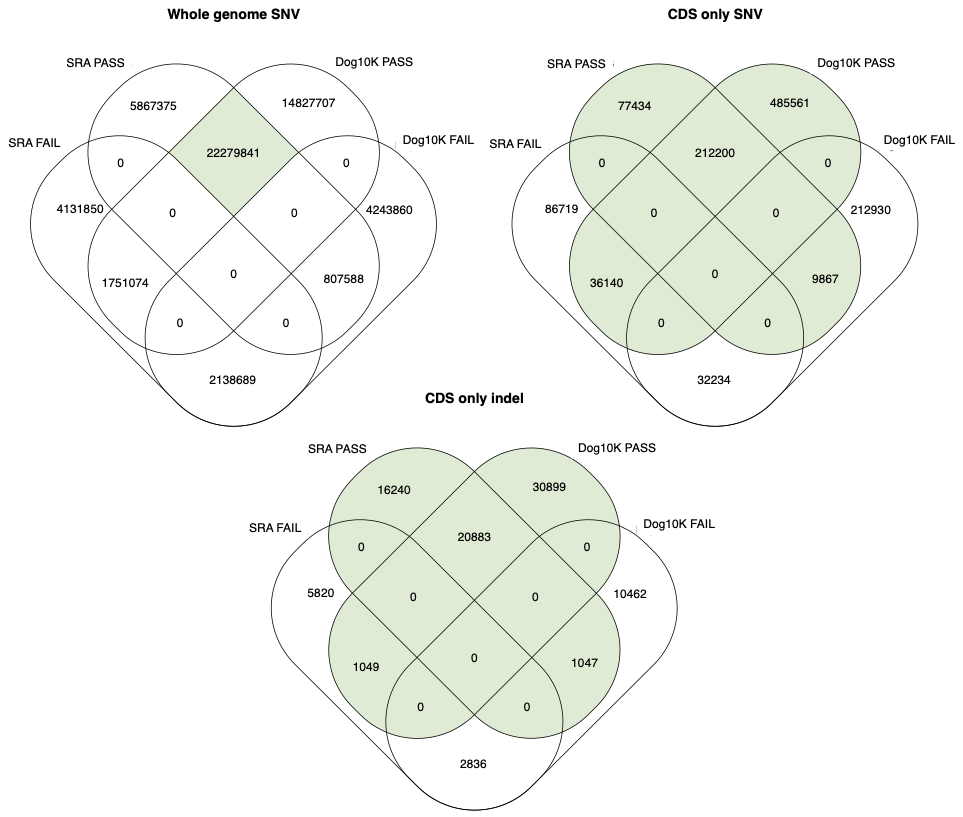

**Supplemental Fig. S11: The number of shared and unique VQSR PASS/FAIL SNV sites from each dataset after sample filtering.** The green shading indicates that for whole genome analyses, only VQSR PASS sites in both datasets were used for further analysis. For coding sequence (CDS) analyses, only sites that are VQSR PASS in either dataset were used for further analysis. Whole genome SNV analyses used a total of 22,279,841 sites, CDS-only SNV analyses used a total of 821,202 sites, and CDS-only indel analyses used a total of 70,118 sites.

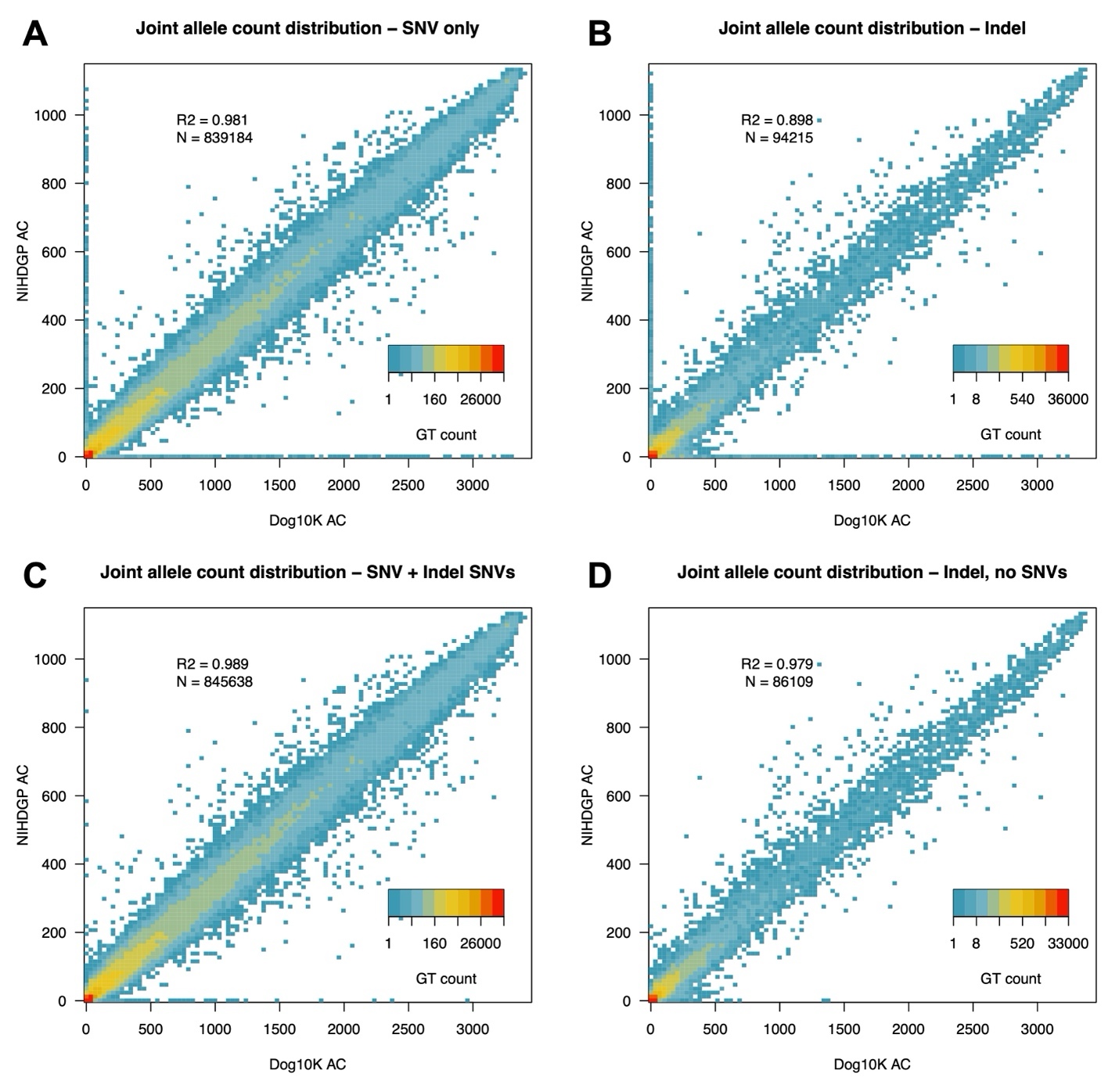

**Supplemental Fig. S12: Allele counts of CDS variants assigned as SNV or indel in the Dog10K and SRA cohorts.** A) Allele counts for variants that were initially identified as SNVs in each cohort. B) Allele counts for variants that were initially identified as indels in each cohort. C) Allele counts of initially identified SNVs and SNVs recovered from the indel set. D) Allele counts of indels after removing SNVs.

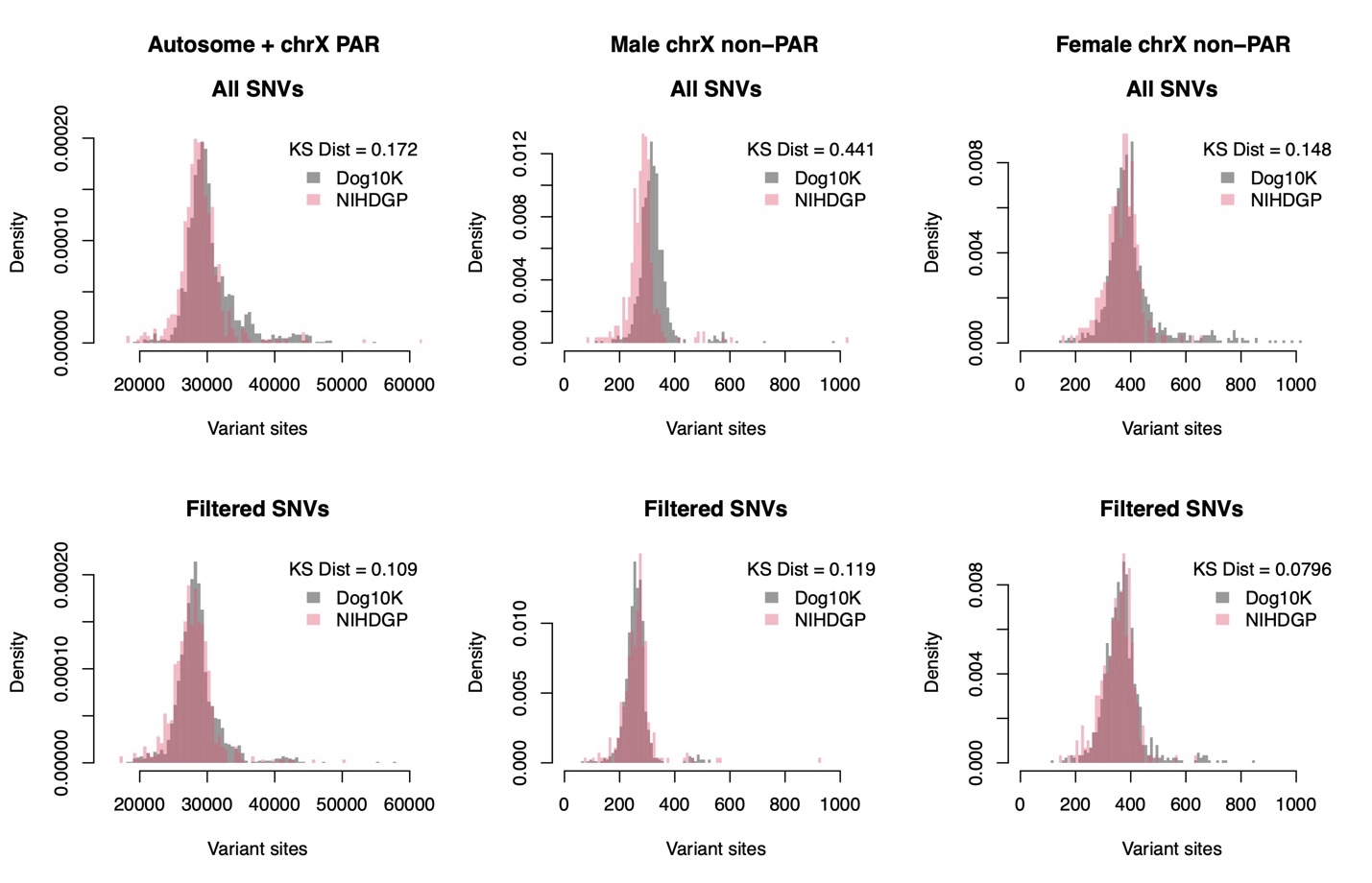

**Supplemental Fig. S13: CDS SNVs per dog before and after genotype filtering.** KS dist represents the Kolmogorov–Smirnov distance of the cumulative densities of each distribution. In each case, genotype filtering leads to both datasets reporting similar numbers of CDS SNVs per dog. Males and females were analyzed separately for the non-PAR region of chrX due to differences in ploidy.

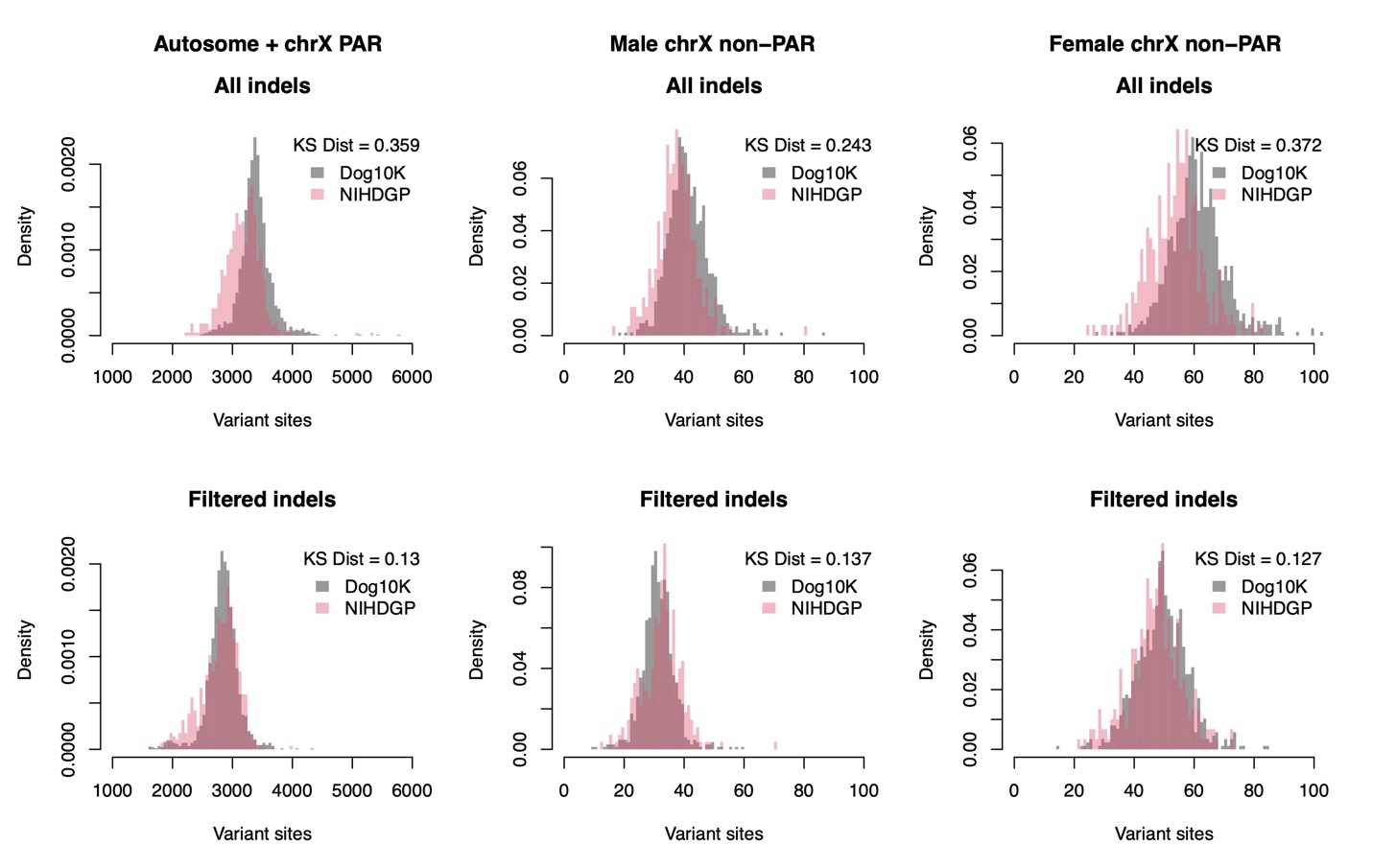

**Supplemental Fig. S14: CDS indels per dog before and after genotype filtering.** KS dist represents the Kolmogorov–Smirnov distance of the cumulative densities of each distribution. In each case, genotype filtering leads to both datasets reporting similar numbers of CDS indels per dog. Males and females were analyzed separately for the non-PAR region of chrX due to differences in ploidy.
